# Acriflavine, a clinically aproved drug, inhibits SARS-CoV-2 and other betacoronaviruses

**DOI:** 10.1101/2021.03.20.436259

**Authors:** Valeria Napolitano, Agnieszka Dabrowska, Kenji Schorpp, André Mourão, Emilia Barreto-Duran, Malgorzata Benedyk, Pawel Botwina, Stefanie Brandner, Mark Bostock, Yuliya Chykunova, Anna Czarna, Grzegorz Dubin, Tony Fröhlich, Michael Hoelscher, Malwina Jedrysik, Alex Matsuda, Katarzyna Owczarek, Magdalena Pachota, Oliver Plettenburg, Jan Potempa, Ina Rothenaigner, Florian Schlauderer, Artur Szczepanski, Kristin Greve-Isdahl Mohn, Bjorn Blomberg, Michael Sattler, Kamyar Hadian, Grzegorz Maria Popowicz, Krzysztof Pyrc

## Abstract

The COVID-19 pandemic caused by SARS-CoV-2 has been socially and economically devastating. Despite an unprecedented research effort, effective therapeutics are still missing to limit severe disease and mortality. Using high-throughput screening, we identified acriflavine as a potent papain-like protease (PL^pro^) inhibitor. NMR titrations and a co-crystal structure confirm that acriflavine blocks the PL^pro^ catalytic pocket in an unexpected binding mode. We show that the drug inhibits viral replication at nanomolar concentration in cellular models, *in vivo* in mice and *ex vivo* in human airway epithelia, with broad range activity against SARS-CoV-2 and other betacoronaviruses. Considering that acriflavine is an inexpensive drug approved in some countries, it may be immediately tested in clinical trials and play an important role during the current pandemic and future outbreaks.

## Introduction

Coronaviruses have been considered a potential threat since 2002, when the severe, acute respiratory syndrome coronavirus (SARS-CoV) emerged in southern China spreading across continents but disappearing shortly thereafter (Drosten et al., 2003; Ksiazek et al., 2003). Ten years later, Middle-East respiratory syndrome coronavirus (MERS-CoV) posed a pandemic threat, but despite high fatality rates, the human-to-human transmission remained limited (Zaki et al., 2012). Despite these warnings, the emergence of the SARS-CoV-2 and subsequent pandemic found healthcare largely unprepared and has paralyzed the modern world in an unprecedented way (Iacobucci, 2020; Ma et al., 2020; Mirzaei et al., 2020).

First vaccines are available and applied already, and further approvals are expected shortly (Parker et al., 2020). However, the time needed for vaccination of the global population, limited availability of vaccines, reluctance to vaccinate, and reduced effectiveness of vaccination against newly emerging variants underline the urgent need for effective antivirals (Saha et al., 2020). The timeline of preclinical to clinical development of novel antivirals, however, is too long for completely new compounds to make a clinical impact during the current pandemic. Repurposing existing drugs with known safety profiles is, therefore, the most efficient and promising option. Initial candidates, unfortunately, did not fulfill expectations (Cao et al., 2020; Horby et al., 2020). Some other leads are still being tested in the clinic, but no convincing proof of efficacy has been provided as yet (Pan et al., 2020; Simonovich et al., 2020). Importantly, it has been proposed that the identification of a set of antivirals against SARS-CoV-2 may provide an opportunity to mimic the strategy effective for HIV-1 - combinatorial treatment against different molecular targets (Lu et al., 2018).

The coronaviral genome encodes several structural and non-structural proteins (Knipe and Howley, 2013), including two cysteine proteases, M^pro^ (nsp5) and PL^pro^ (nsp3), essential for the virus replication (Shamsi et al., 2021). While extensive actions to identify inhibitors of M^pro^ have been undertaken (Capasso et al., 2020), only scarce information is available for the PL^pro^. The latter protease is essential for viral protein maturation and has also been attributed to other functions, including type I interferon response attenuation, making it a viable therapeutic target (Shin et al., 2020).

Here, we discovered acriflavine (ACF) as an effective inhibitor of SARS-CoV-2. Acriflavine (ACF) is a mixture of trypaflavines (3,6-diamino-10-methylacridinium chloride and 3,6-diamino-3-methylacridinium chloride) and proflavine (3,6-diaminoacridine). ACF has been used systemically against sleeping sickness, urinary tract infections, and gonorrhea and has been suggested effective in a number of other indications (Dana et al., 2014; Funatsuki et al., 1997; Kawai and Yamagishi, 2009; Manchester et al., 2013; Mathé, 2000; Pépin et al., 2017; Persinoti et al., 2014; Tripathi et al., 2006). ACF was tested in a clinical trial against HIV where up to 100 mg daily was administered over the course of several months. Despite the chemical structure suggesting possible DNA intercalation and liver toxicity, no adverse effects were observed in these studies (Mathé et al., 1996; Mathé et al., 1998). ACF has been approved in the past and was present in European pharmacopeias for systemic use (Browning, 1943, 1967; Mathé et al., 1998; Wainwright, 2001). In some countries, ACF is currently available as an over-the-counter medicine (e.g., Brazil – EMS Cystex) against urinary tract infections. Its safety and efficacy were tested in a recently reconfirmed in a clinical trial in Brazil (Gama CRB, 2020; NCT03379389, 2020). In other countries, ACF is used as a content of mouthwash advised for children (“Michino” gargle in Japan) (Lee et al., 2009). The extensive available studies on the long-term systemic administration of large doses of ACF have not revealed any major side effects, suggesting that it could be employed alone or in combination with other drugs (Nehme et al., 2020).

Our data demonstrate that ACF is a potent inhibitor of SARS-CoV-2 PL^pro^, opening a promising therapeutic approach against COVID-19. The mode of action is confirmed by structural characterization with X-ray crystallography and NMR, demonstrating that ACF specifically inhibits the active site of the enzyme with an unprecedented binding mode. We show that ACF blocks the virus infection *in vitro*. Importantly, the viral infection is also suppressed *ex vivo* in human airway epithelium (HAE) cultures, which recapitulate the fully-differentiated human airway epithelium, as well as in an *in vivo* mouse model. Finally, we provide evidence for ACF’s antiviral effect against other betacoronaviruses, making this a potentially important therapeutic against future pandemics.

## Results

### Identification and validation of PL^pro^ repurposed inhibitors

ACF was identified in a high-throughput screening approach for inhibitors of PL^pro^. We established a protease assay using a fluorogenic substrate. The overall scheme of this study is presented in **Figure 1A**. Based on the screening of a library consisting of 5,632 small molecule compounds, which are either approved or are/have been at various stages of clinical testing (Corsello et al., 2017), we selected 11 compounds for validation experiments (**Supplementary Table 1**; **Supplementary Figure S1**). Acriflavine (ACF) was the most active compound with a dose-dependent PL^pro^ inhibition with an IC_50_ of 1.66 μM (**Figure 1B**). Next, we tested the effect of ACF on PL^pro^ enzymatic activity using the ISG15-AMC probe as substrate (Klemm et al., 2020; Shin et al., 2020). As shown previously, PL^pro^ cleaves ISG15-AMC significantly faster compared to RLRGG-AMC. Nonetheless, the IC_50_ of ACF with ISG15-AMC (1.46 μM) (**Figure 1B, Supplementary Figure S2B;**) was comparable to the IC_50_ determined with the RLRGG-AMC substrate. To eliminate possible fluorescence artifacts, we conducted gel-based de-ubiquitination assays to confirm our results. We incubated K48 tri-ubiquitin with PL^pro^ and took samples at different time points to analyze the cleavage by Western Blot. PL^pro^ efficiently cleaved K48 tri-ubiquitin to di-ubiquitin (**Figure 1C**) as previously shown (Klemm et al., 2020).

**Figure 1.**
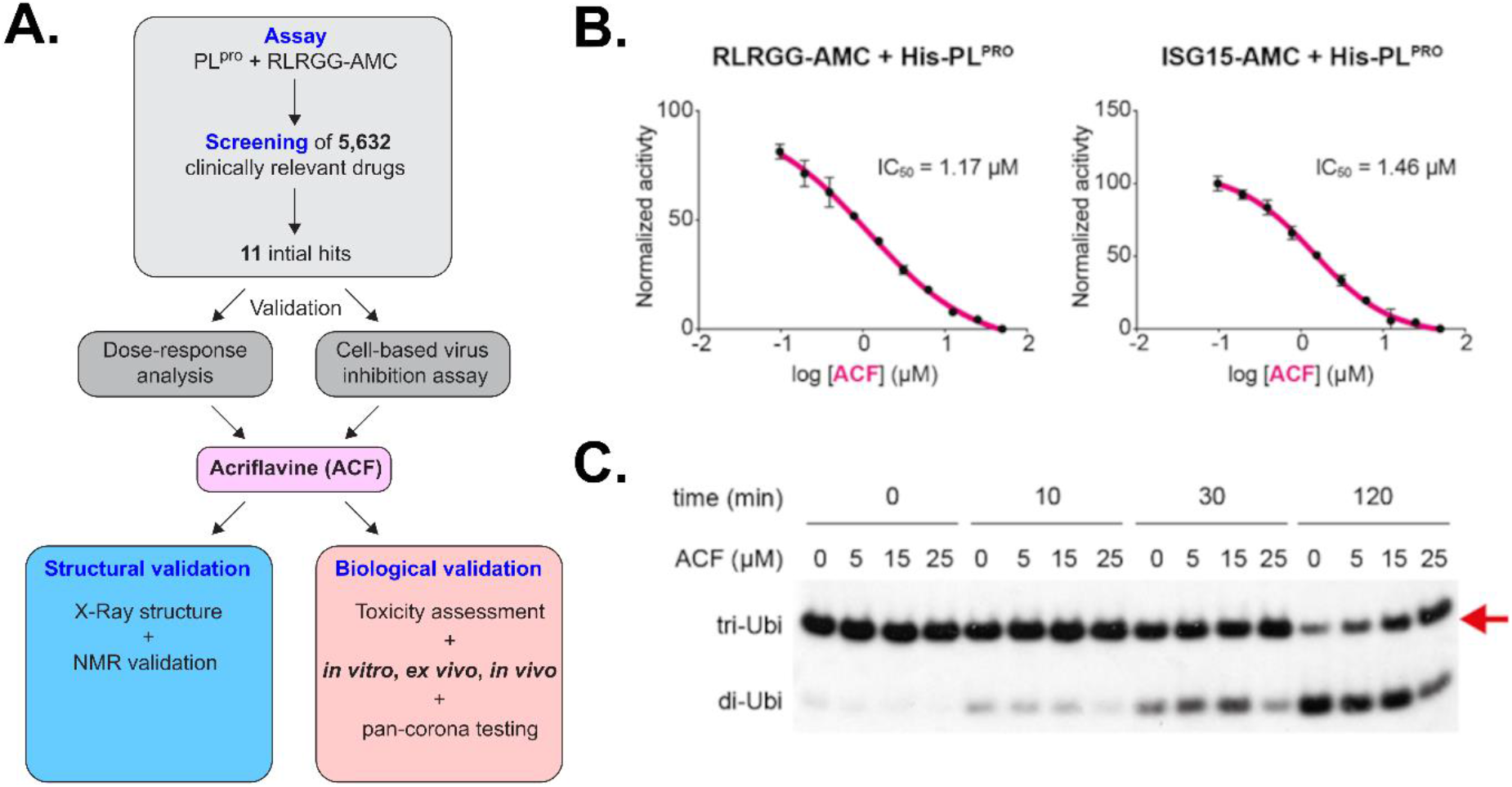
Identification of PL^pro^ inhibitors. ACF as a confirmed hit. (**A.**) General scheme of the study. (**B.**) Enzymatic activity inhibition assay using RLRGG-AMC (left) or ISG15-AMC (right) as substrates for PL^pro^ performed in technical triplicates. The vertical axis shows fluorescence intensity (rising, as the substrate is proteolytically cleaved) as a function of time. (**C.**) Time-course analysis of tri-ubiquitin K48-linked (2 μM) hydrolysis using 100 nM PL^pro^ in the presence of different ACF concentrations.

Next, PL^pro^ was incubated with either DMSO or 5, 15, 25 μM of ACF before adding K48 tri-ubiquitin chains to the reaction mixture. Also, in this independent assay format, ACF reduced the protease activity in a dose-dependent manner (**Figure 1C**), thereby confirming that ACF is a specific PL^pro^ inhibitor.

To evaluate the specificity of ACF we performed an enzymatic digestion assay using fluorescent M^pro^ substrate. This experiment shows that ACF is a very weak M^pro^ inhibitor with less than 50% inhibition at 100 μM of ACF concentration (**Supplementary Figure S3**). Therefore, we conclude that M^pro^ inhibition by acriflavine is not a significant force in reducing SARS-CoV-2 replication.

### Structural analysis of SARS-CoV-2 PL^pro^ in complex with proflavine

In order to confirm that ACF binds directly to PL^pro^ in solution we recorded 2D ^1^H,^15^N TROSY NMR spectra of recombinant ^15^N-labeled PL^pro^ in the absence and presence of ACF. The spectra of PL^pro^ are consistent with a globular folded, monomeric protein in solution (**Figure 2A**). The addition of ACF induces chemical shift changes for a number of amide signals while not affecting the remaining bulk of the resonances. This confirms that the overall fold and monomeric state of the protein are not affected and demonstrates that ACF binds to a spatially localized site in PL^pro^.

**Figure 2.**
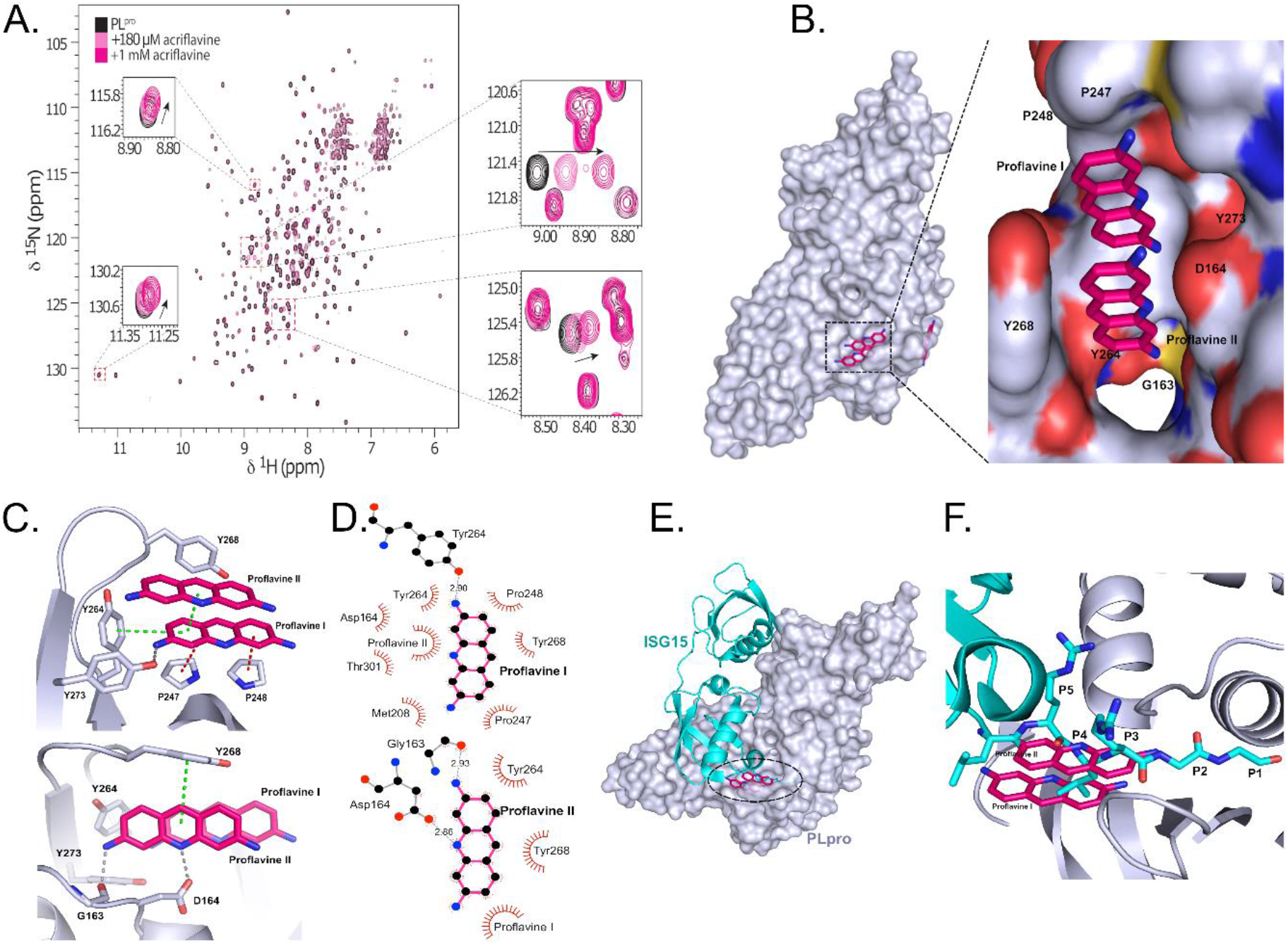
Structural details of PL^pro^ inhibition by proflavine. **(A.)** Overall crystal structure of PL^pro^-proflavine complex at 2.7 Å. The magnified fragment shows two proflavine molecules inside the substrate recognition cleft of PL^pro^. **(B.)** Intermolecular interaction between PL^pro^ and proflavine molecules. Two π-stacked molecules form a network of hydrogen bonds, π-π interactions, and hydrophobic contacts with PL^pro^ **(C.)** Molecular interaction details for the two proflavine molecules. **(D. E.)** Overlay of the ISG15-PL^pro^ structure and PL^pro^-proflavine structure. Proflavines are bound in the same place as the C-terminal end of the ISG15 substrate and mimic polar and lipophilic interactions with the native substrate. **(F.)** 1^H^,15^N^ heteronuclear correlation NMR spectra of PL^pro^ titrated with ACF demonstrate the localized binding and unaltered monomeric state of the protein.

To understand the molecular basis of the SARS-CoV2-PL^pro^ inhibition by acriflavine, we determined the X-ray crystal structure of the protease in complex with proflavine, one of the principal components of ACF. Analysis of the electron density map shows that two molecules of proflavine are π-π stacked to each other and accommodate the S3-S5 pockets of the PL^pro^ substrate recognition cleft, which is defined by the loop connecting the helices α3 and α4 and the so-called “blocking loop” BL2 (**Figure 2B**) (Ratia et al., 2006). Comparison with the apo structure of PL^pro^ (PDB code: 7D47) revealed that, although the overall structure is well preserved, the BL2 loop undergoes a large conformational change upon binding of the ligand. In particular, the side chain of Tyr268 rotates by around 57° inwards towards the substrate recognition cleft, and the BL2 loop moves by 2 Å in the same direction, narrowing the substrate-binding cleft (**Supplementary Figure S4**).

Several PL^pro^ residues are involved in ligand binding. One of the molecules termed proflavine-I occupies the S4 pocket (**Figure 2B, C, D**). The side chain Tyr273 is involved in a hydrogen bond (2.9 Å) with the primary amine group at position 3 of proflavine I, which sits at the bottom of the substrate-binding cleft, where it forms a T-stacked CH/π interaction with Pro247 (3.9 Å). A parallel stacking CH/π interaction with Pro248 (4.6 Å) and T-shaped π-π stacking with the aromatic side chain of Tyr264 is engaged in a T-shaped π-π stacking interaction (3.5 Å) further stabilize the interaction with proflavine I. The second molecule, termed proflavine-II, shows π-π stacking at 3.5 Å with proflavine-I and occupies the S3 and S5 pockets (**Figure 2D, E**). Gly163 and Asp164 form hydrogen bonds with the primary amine group at position 3 (2.9 Å) and the imino group of the acridine moiety (2.9 Å), respectively. In addition, Tyr268 forms a T-shaped π-π stacked interaction (5.1 Å) with proflavine-II. Overall, this provides a unique binding mode, where two proflavines, tightly π-π stacked, cooperate in blocking the substrate pocket. Importantly, both molecules are required for the inhibition.

Recently, Shin *et al.* reported the crystal structure of SARS-CoV2-PL^pro^ in complex with ISG15 (interferon-induced gene 15) bearing the RLRGG recognition motif at the C-terminus (Shin et al., 2020). Comparison with our structure shows that the proflavine-II molecule, which occupies the S3 and S5 pockets, overlaps very well with the backbone of Arg151 and Arg153 in the RLRGG peptide in positions P3 and P5, respectively (**Figure 2E, F**). The side chain of Leu152 in position P4 points exactly towards the other molecule of proflavine in the S4 pocket. All the major interactions of the substrate with PL^pro^ residues are recapitulated by the inhibitor. Mechanistically, these observations indicate that proflavine inhibits SARS-CoV2-PL^pro^ by limiting access to the substrate-binding site. Moreover, the fact that the imino group of the proflavine-I molecule in the S4 pocket is not involved in any polar interactions suggests that the methylated (acriflavine) form is preferred at this position due to lack of desolvation penalty. The amino group at position 3 of proflavine-II is located in a similar position as the amide nitrogen of the RLRGG glycine P2 in the P2 pocket. Side-methylated proflavine, which is also present in commercial acriflavine preparations (see Supplementary Analysis), would, therefore, mimic the P2 amino acid. These two expected interactions of commercial acriflavine explain why it is a significantly more potent inhibitor than pure proflavine. Intriguingly, a unique and important feature of the commercial acriflavine mixtures is that they contain the combination of methylated proflavines to optimally block the PL^pro^ active site.

### ACF activity in SARS-CoV-2 infected cell culture

For cellular validation of ACF activity, two SARS-CoV-2 infection cell culture models were used: Vero cells, which are a broadly used simian model, and human A549^ACE2+^ cells overexpressing the ACE2 receptor (Milewska et al., 2020). First, the cytotoxicity of ACF was evaluated on cell lines (A549^ACE2+^, Vero, HCT-8, CRFK) as well as primary human fibroblasts (**Figure 3A, Supplementary Figure S5**). The cellular cytotoxicity CC_50_ values were determined in both cell types to be 3.1 μM for A549^ACE2+^, 3.4 μM for Vero, 2.1 μM for HCT-8, and 12 μM for primary human fibroblasts. Interestingly, ACF shows lower cytotoxicity in primary cells compared to transformed cell lines, which may reflect the previously described antineoplastic activity of ACF (Lee et al., 2014). Secondly, dose-response experiments were carried out in A549^ACE2+^ and Vero cells (**Figure 3B**). The IC_50_ values determined for ACF are 86 nM and 64 nM for A549^ACE2+^ cells and Vero cells, respectively. Accordingly, the selectivity index (SI) values for A549^ACE2+^ and Vero cells are 36 and 53, thus representing a potent inhibitor of SARS-CoV-2 virus replication with a favorable cytotoxicity profile. Notably, ACF inhibition at 400 nM is significantly more effective than inhibition by remdesivir at 25-fold higher concentration in these assays (**Figure 3B**).

**Figure 3.**
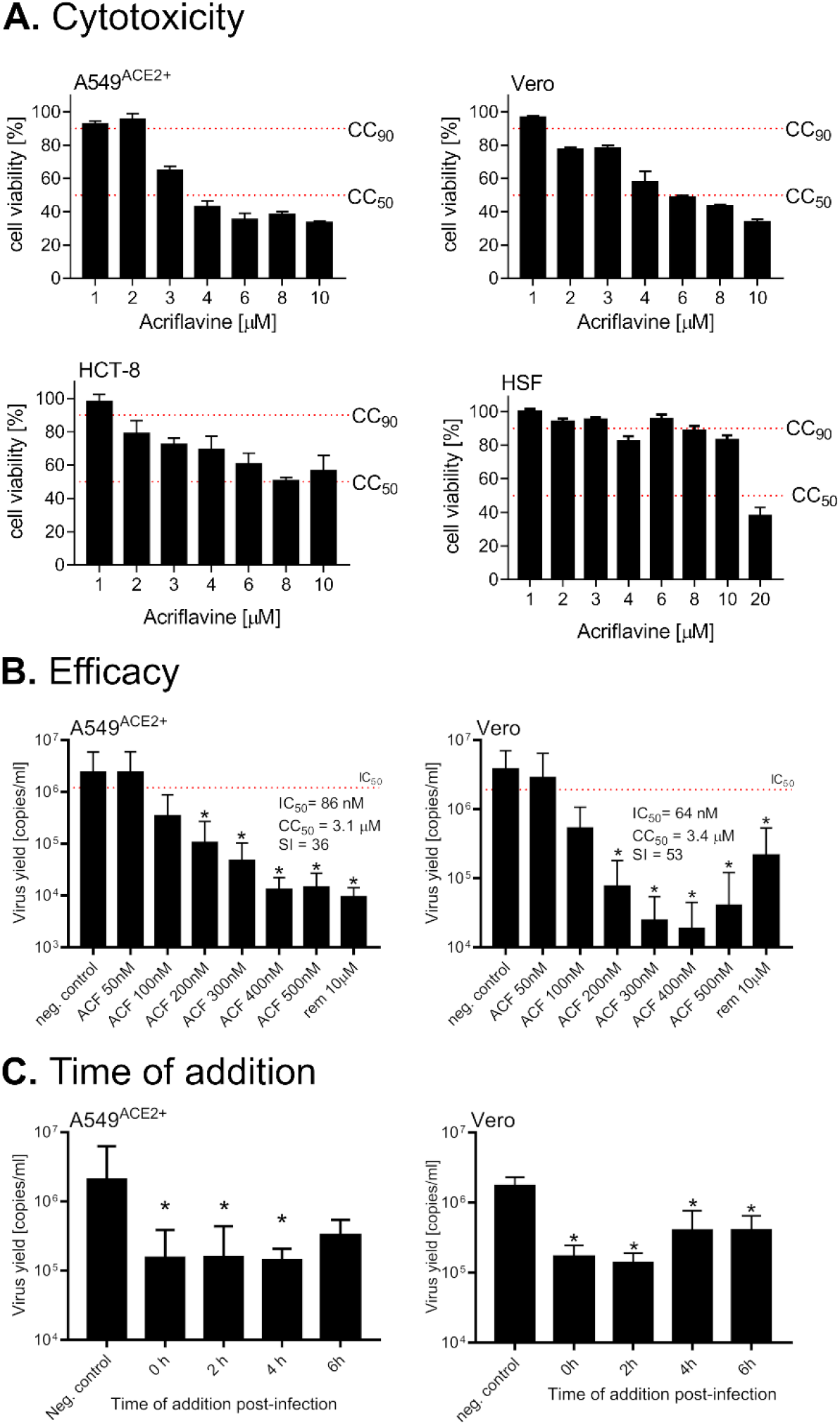
ACF inhibits SARS-CoV-2 replication *in vitro*. (**A.**) The cytotoxicity of ACF in four cell culture systems employing cell lines (A549ACE2+, Vero cells, HCT-8 cells) and primary human skin fibroblasts (HSF). (**B.**) Inhibition of virus replication by ACF in A549^ACE2+^ (left panel) and Vero (right panel) cells. The figure shows RT-qPCR analysis of cell culture supernatants infected with SARS-CoV-2 at 1600 TCID_50_/ml 24 h post-infection. (**C.**) Time of addition study. The inhibition of virus replication in A549^ACE2+^ cells (left panel) or Vero cells (right panel) by ACF added at different times post-infection (time denoted on the x-axis). The figure shows RT-qPCR analysis of cell culture supernatants infected with SARS-CoV-2 at 1600 TCID_50_/ml) 24 h post-infection. All experiments were performed at least in 3 biological repetitions, each in triplicate. The results are presented as average values with standard deviations (error bars). An asterisk indicates values that are significantly different from the control (*p* < 0.05).

Next, we carried out time-of-addition experiments, where cells (A549^ACE2+^ and Vero cells) were first infected with the virus, and the ACF was added after 0, 2, 4, or 6 h. The results show that virus inhibition was maintained, even when the compound was not present during the early stages of infection. Consequently, we conclude that SARS-CoV-2 is mainly inhibited during the replication phase, and ACF does not affect viral entry (**Figure 3C**).

Inhibition of virus replication by ACF was further confirmed using confocal microscopy. 24 h and 48 h post-infection in the presence or absence of 500 nM ACF or 10 μM remdesivir, confocal microscopy images showed nearly complete elimination of SARS-CoV-2 when compared to untreated cells. While virus elimination is also observed for remdesivir, this is only seen at more than 20-fold higher concentrations compared to ACF (**Figure 4A**).

**Figure 4.**
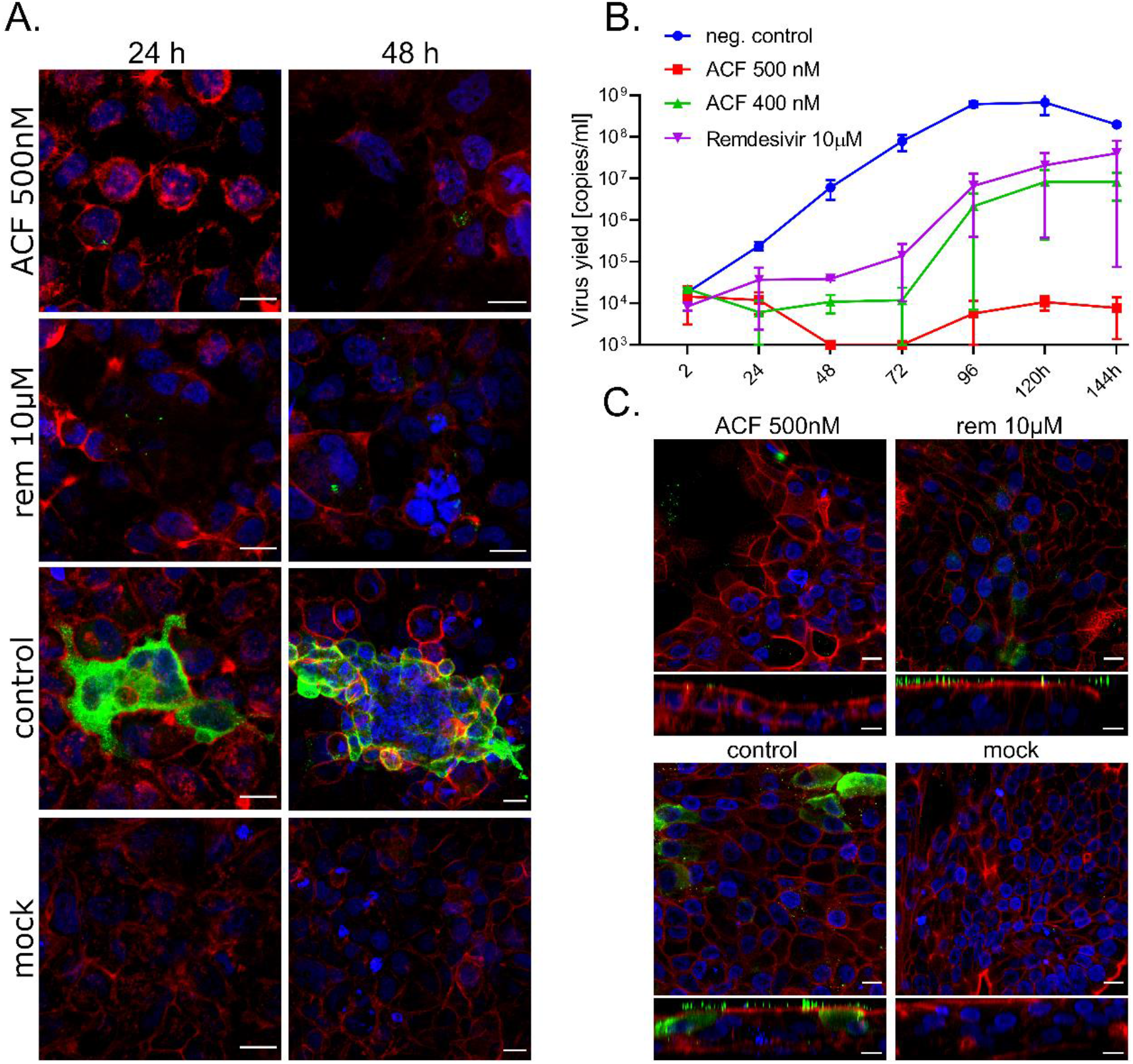
ACF blocks SARS-CoV-2 replication *in vitro* **and** *ex vivo*. **A.** The inhibition of virus replication by ACF in A549^ACE2+^ cells visualized with confocal microscopy. Cells were infected with SARS-CoV-2 in the presence of 500 nM acriflavine, 10 μM remdesivir, or vehicle control (control) for 24 h and 48 h. Cell nuclei are denoted in blue, actin is denoted in red, and SARS-CoV-2 N protein is denoted in green. Each image represents maximum projection of 5 μm section. Scale bar = 20 μm. **B.** Antiviral activity of ACF against SARS-CoV-2 in human airway epithelium (MucilAir). The figure shows RT-qPCR analysis of HAE culture supernatants infected with SARS-CoV-2. Remdesivir and PBS were used as controls. The assay was performed at least in duplicate, and median values with range are presented. Two-way ANOVA analysis with Dunnett’s post-hoc test indicated that ACF and REM significantly inhibit virus yields during infection course compared to untreated control. (**C.**) Confocal analysis of infected HAE cultures. Cells were infected with the SARS-CoV-2 in the presence of 500 nM acriflavine, 10 μM remdesivir, or vehicle control (control). Mock images illustrate non-infected cells. On day 6 p.i. cells were fixed and immunostained. Cell nuclei are denoted in blue, actin in red and SARS-CoV-2 N-protein in green. Each image represents maximum projection of 3 μm section. Scale bar = 10 μm.

### *Ex vivo* inhibition of SARS-CoV-2 infection in HAE cultures

The antiviral activity of ACF was analyzed in the HAE *ex vivo* model. Two different concentrations were evaluated (400 nM and 500 nM); PBS and remdesivir were used as controls. The results (**Figure 4B, Supplementary Figure S6**) show inhibition of SARS-CoV-2 replication in the presence of ACF and remdesivir in the HAE *ex vivo* model. A significantly lower viral yield was detected in the cultures treated with ACF in comparison to the PBS control. 500 nM ACF treated HAE showed higher inhibition of virus replication than the positive control remdesivir at 10 μM concentration after 144 h of infection. At time points of 48 and 72 hours of infection, the virus was under the detection limits in the ACF 500 nM treated sample. Importantly, while virus titers upon 10 μM remdesivir treatment relapsed after 144 hours to control levels, 500 nM ACF treatment kept the virus at low titers.

After 6 days of infection in the presence of 500 nM ACF, confocal microscopy images showed complete inhibition of SARS-CoV-2 replication. While inhibition was also noted for remdesivir. However, the virus particles in remdesivir treated cells were still present on the cell surface, suggesting that a detectable level of virions was still produced. Therefore, the inhibition by 10 μM remdesivir was inferior compared to 500 nM ACF (**Figure 4C**).

### Pan-coronavirus activity of ACF

To verify whether ACF may be used as a broadband anticoronaviral drug, its activity was tested against betacoronaviruses other than SARS-CoV-2, *i.e.,* MERS-CoV and HCoV-OC43, as well as alphacoronaviruses, *i.e.,* HCoV-NL63 and feline infectious peritonitis virus (FIPV). ACF efficiently inhibited MERS-CoV (IC_50_ = 21 nM, SI = 162) and HCoV-OC43 (IC_50_ = 105 nM, SI = 27). Interestingly, no effect on the replication of tested alphacoronaviruses was observed (**Figure 5)**. These data reveal that ACF can be used to inhibit betacoronaviruses in a broader sense and could represent a drug for future outbreaks.

**Figure 5.**
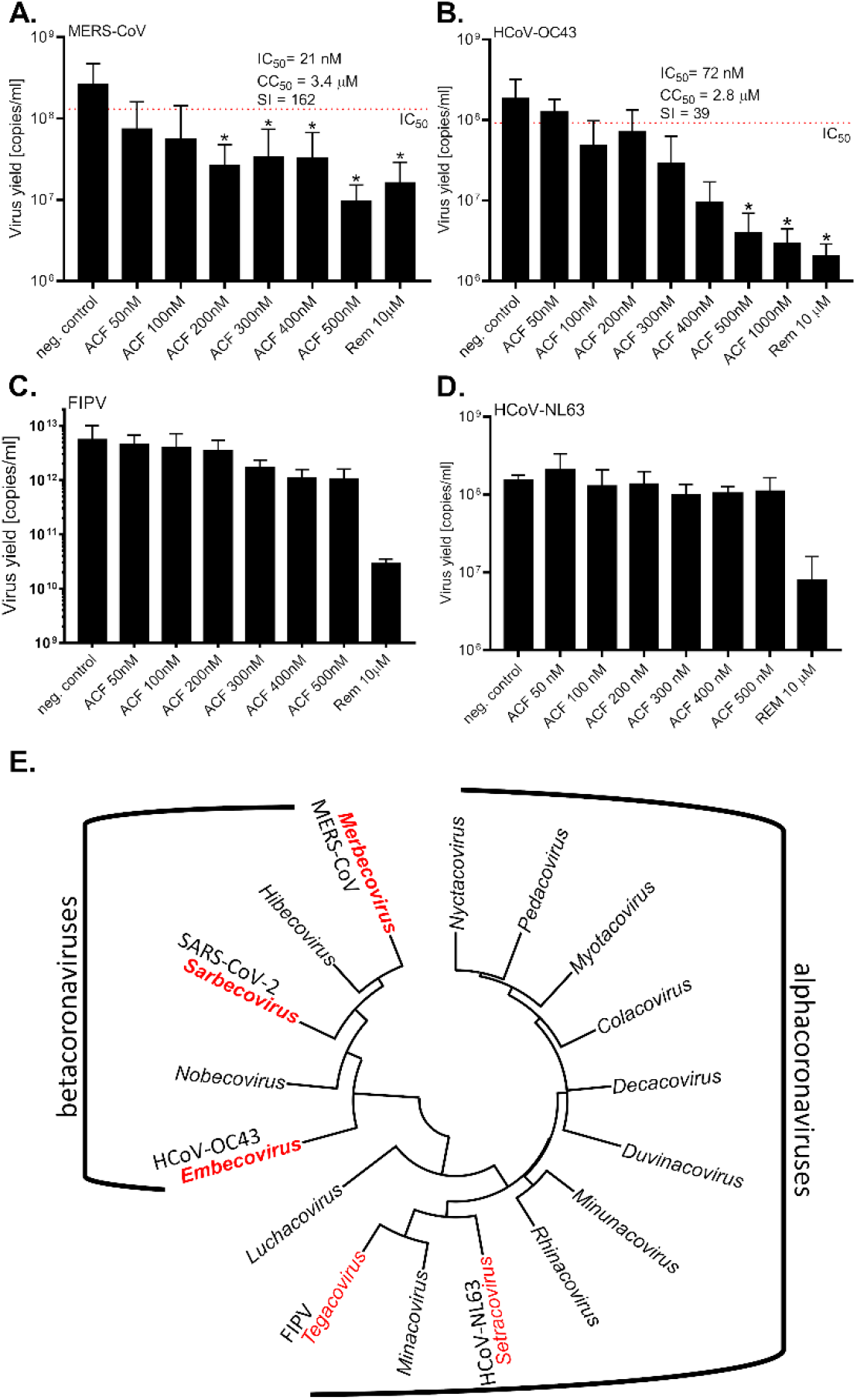
ACF is a broad-spectrum inhibitor of betacoronaviruses. Replication of (**A.**) MERS-CoV, (**B.**) HCoV-OC43, (**C.**) HCoV-NL63 and (**D.**) FIPV *in vitro* in the presence or absence of inhibitors, as assessed with RT-qPCR on cell culture supernatants. A single round of infection was recorded (24 h). All experiments were performed at least in 2 biological repetitions, each in triplicate. The results are presented as average values with standard deviations (error bars). An asterisk indicates values that are significantly different from the non-treated control (*p* < 0.05). ACF: acriflavine, rem: remdesivir. (**E.**) Phylogenetic tree of alpha and betacoronaviruses. Three different genera of betacoronaviruses were used in the study as representants and are denoted in red. Representative viruses used in this study are denoted in red.

### Effect of ACF in an *in vivo* model of infection

To confirm that ACF can block virus replication *in vivo* we used an infection model based on K18-ACE2 mice. The experiment groups consisted of infection control receiving no treatment, ACF administered orally (*p.o.*; 100 mg/kg daily), two groups receiving ACF intramuscularly (*i.m.*; 5 mg/kg or 15 mg/kg body weight twice a day), and a positive control group receiving remdesivir (*i.m.*; 25 mg/kg per day). We chose a high dose of oral administration to alleviate low bioavailability and relatively fast elimination of the drug. The administration of the ACF and remdesivir started one day prior to infection with SARS-CoV-2 virus. The treatment was continued for six consecutive days.

In all groups, the animals survived the infection until the last day, and no apparent signs of the disease were noted, including lack of change in the body weight. There were also no drug-related adverse effects. Analysis of the virus replication levels by RT-qPCR on day 6 in the untreated group clearly shows that the virus replicated in lungs and brains. Treatment with acriflavine (both p.o. and i.m.) almost completely blocked the infection in the brain (**Figure 6A**), while the positive control remdesivir only decreased the viral loads by 2-3 orders of magnitude. The infection levels were less pronounced in the lungs, but suppressed replication of the virus was also recorded for treatment groups, with the most pronounced effect (1-2 orders of magnitude reduction) observed for the p.o. administration of ACF (**Figure 6B**).

**Figure 6.**
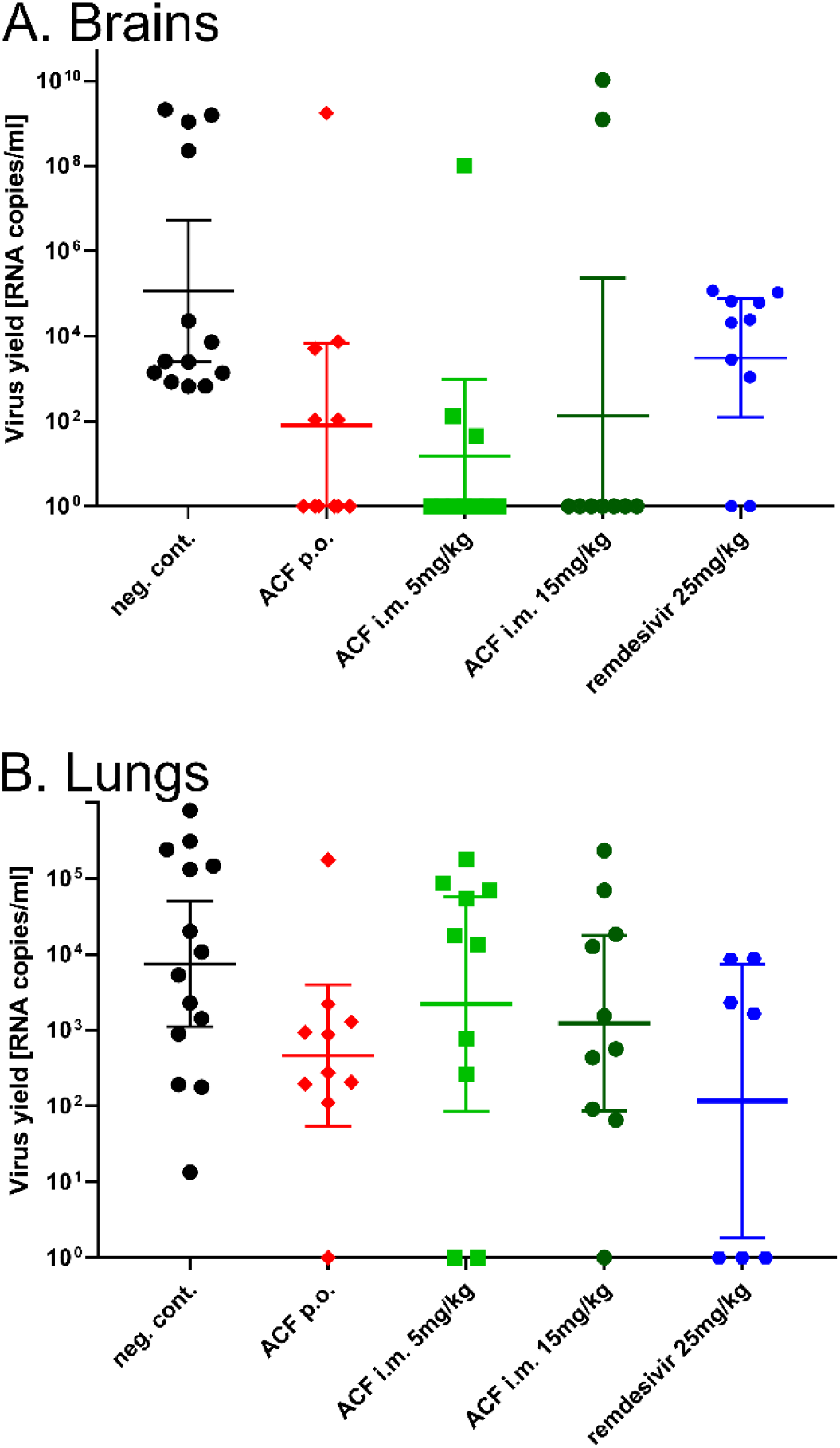
Inhibition of virus replication by ACF *in vivo*. Each group consisted of 10 animals. Oral and intramuscular delivery were tested. Remdesivir was used as positive control. Virus replication in different organs was assessed using the RT-qPCR. Strong reduction of virus level is visible in the brain of treated groups. ACF caused a marked reduction of virus load in the lungs when administered orally. Data are presented as geometric mean with 95% confidence intervals.

## Discussion

SARS CoV-2 is the cause of the most devastating global pandemic since the 1918 influenza. Despite the large commitment, early efforts failed to deliver effective antivirals to control the infection. Drugs inhibiting virus replication should be given early during the infection, but the required early diagnosis and the availability and costs involved render this challenging. A cheap, easily accessible drug-like ACF might offer the required early availability and accessibility. Moreover, compounds of limited efficacy in monotherapy may serve as part of a combination therapy strategy (Lu et al., 2018), as the synergy between the compounds targeting different molecular targets may be favorable.

Here, we focused on one of the viral proteases, PL^pro^, which is indispensable for SARS-CoV-2 replication, progeny production, and attenuation of the host’s immune responses (Knipe and Howley, 2013; Shin et al., 2020). PL^pro^ modulates the ISGylation pathway, which acts as a viral defense mechanism to inhibit normal protein translation. ISGylation modifies protein kinase R, so that newly synthesized (viral) proteins are tagged as dangerous (Villarroya-Beltri et al., 2017). Inhibition of PL^pro^ will not only interfere with virus protein maturation, but additionally will activate the host cellular defense mechanisms to fight SARS-CoV-2 infection.

We show that ACF is a potent, specific inhibitor of PL^pro^, with IC_50_ values in the low nanomolar range in all tested cell models. The activity is promising and by far exceeds the reported activities for GRL-0617 (Shin et al., 2020) and rac5c (Klemm et al., 2020), both active at the micromolar range. Structural analysis revealed that the inhibitor blocks the catalytic site of the PL^pro^. An unexpected and highly unusual mechanism of inhibition is observed, where two ACF molecules bind in the enzyme’s substrate binding site. The crystal structure and validation of binding in solution using NMR confirm the mode of action of ACF. Together with the high potency in cellular models these findings identify ACF as a highly attractive repurposing molecule to inhibit SARS-CoV-2 replication.

Despite a relatively short half-life for ACF *in vivo* (Song et al., 2005) we have observed several orders of magnitude reduction of viral load in a humanized mice infection model. This suggests that with the development of the correct treatment regime, the therapeutic effect of ACF can be extended in humans.

Notably, in addition to SARS-CoV-2, the antiviral effect of ACF was observed for other betacoronaviruses: MERS-CoV, and HCoV-OC43, but not for other genera of the *Coronaviridae* family (alphacoronaviruses). We believe that these data indicate broad-spectrum activity of ACF against betacoronaviruses, with the potential to target future zoonotic coronaviruses. This means ACF has therapeutic benefits not only in the current crisis but can also become an important component of the arsenal against future pandemics.

The unique and specific inhibition of SARS-CoV-2 PL^pro^ and associated inhibition of virus replication in the cell culture, *ex vivo* and *in vivo* renders ACF an attractive compound for evaluation in the treatment of COVID-19. It has to be considered, though, that the aromatic acridine rings are linked with DNA intercalating properties and, consequently, with mutagenic potential. On the other hand, a substantial amount of long-term clinical data is available for systematic applications (Gama CRB, 2020; Mathé et al., 1996; Mathé et al., 1998), thus alleviating concerns related to the potential mutagenic effects of short-term antiviral therapy. Importantly, the antiviral effects are observed at concentrations of two to three orders of magnitude lower than those used in experiments assessing *in vitro* mutagenicity of ACF (Obstoy et al., 2015; Rees et al., 1989).

In conclusion, we have identified a potent small-molecule inhibitor of the coronaviral PL^pro^ that is registered and marketed as an inexpensive over-the-counter drug. The low-nanomolar IC_50_ and good selectivity index, combined with the profound inhibition of the viral replication in all tested models, suggest that the drug may be effective in monotherapy or in combination therapy with, e.g., polymerase inhibitors. A possible limitation for human application might be that currently used therapeutic doses, which have been proven to be safe, will not lead to high enough human systemic exposures. As there are no human pharmacokinetic data available yet, systematic PK studies proving that exposures similar to these that have been effective in mice can also be achieved in humans and are well tolerated. Similarly, like other antiviral drugs, ACF may be administered early during the infection to enable the maximum effect. Specifically, an elderly patient population may significantly benefit from a new antiviral treatment option, as mortality in the age group 70+ is significantly increased compared to younger groups. In addition, the antiviral effect against other betacoronaviruses argues that further research into the therapeutic potential of ACF may provide protection against future zoonotic infections of betacoronaviruses. The unique mode of inhibition of ACF will also serve as a blueprint to develop new, improved drugs with a broadband activity using structure-based drug development to prepare for future coronavirus outbreaks.

## Supporting information

6. ACF supplementary analysis

## Acknowledgments

This work was supported by the subsidy from the Polish Ministry of Science and Higher Education for the research on the SARS-CoV-2 and a grant from the National Science Center UMO-2017/27/B/NZ6/02488 to KP.

## Author Contributions

Investigation: VN, KS, FS, AM, AD, EBD, MB, PB, YC, AC, GD, KO, MP, JP, AS, KGIM, BB

Conceptualization, Funding acquisition, Writing – original draft, Supervision: GP, KP, MS, KH, MH

Writing – review & editing: all authors

## Data and materials availability

All data are available in the main text or the supplementary materials.

