## Supplementary Materials and Methods for "Acriflavine, a clinically aproved drug, inhibits SARS-CoV-2 and other betacoronaviruses"

#### Plasmids and proteins

Gene encoding the $PL^{pro}$ (SARS-CoV-2 nsp3 - 746-1060) was synthesized by Integrated DNA Technologies (IDT-Coralville, USA) optimized for *E. coli* expression and cloned using the Slice method (*1*) into several vectors, including petM11, petM13 with a C-terminal $His_6$, pET22b-CPD and a new expression vector named petM5a, a modified petM11 vector. The petM5a vector contains an insertion of optimized codon sequence after the starting codon ATG, GTG AAG TAT CAA AAG, based on Verma *et al.* (*2*), and followed by the N- terminal $His_6$ tag, and by a TEV protease site. The vector map can be found at HMGU-PEPF website. The plasmids were transformed into *E. coli* BL21 (DE3), and the transformed cells were cultured at 37°C in terrific broth (TB) media containing 100 mg/l kanamycin. After the $OD_{600}$ reached 2, the culture was cooled to 18°C and supplemented with 0.25 mM IPTG. The vector petM5a yielded more soluble protein and was used for all protein expression. For isotopically labeled protein, pre-culture was grown in M9 minimal media followed by inoculation ($OD_{600}$ 0.05) into 1 l of $D_2O$ M9 minimal media supplemented with $^{15}$N-ammoniun chloride. After $OD_{600}$ reached 0.8, the culture was cooled to 18°C and supplemented with 0.25 mM IPTG. After overnight induction, the cells were harvested by centrifugation, and the pellets were resuspended in lysis buffer (20 mM Tris-HCl, pH 8.5, 350 mM NaCl, 10% glycerol, 10 mM imidazole, 5 mM beta-mercaptoethanol) and sonicated at 4°C. The insoluble material was removed by centrifugation at 24,000 rpm. The fusion protein was first purified by Ni-NTA affinity chromatography, the supernatant was applied to nickel resin and washed with 10 times the column volume of lysis buffer followed by a wash step of lysis buffer supplemented with 20 mM imidazole. The protein was eluted with 3 column volumes of a buffer containing high imidazole concentration (20 mM Tris-HCl, pH 8.5, 350 mM NaCl, 5% glycerol, 350 mM imidazole, 5 mM beta-mercaptoethanol). 1 mg of TEV protease was added, and the solution was dialyzed overnight at 4°C against a buffer containing low imidazole concentration, 20 mM Tris-HCl, pH 8.5, 150 mM NaCl, 5% glycerol, 10 mM imidazole, 1 mM beta mercaptoethanol. For the final step, the protein was applied to a size exclusion chromatography column, High load S75 (GE-Helathcare, Chigaco, USA), pre-equilibrated with the final buffer, 20 mM Tris pH 8.0, 40 mM NaCl and 2 mM DTT. The purity of the protein was accessed by SDS-PAGE gel.

$M^{pro}$ construct comprising amino acids 3264-3569 of SARS-CoV-2 polyprotein 1ab, N-terminal 6xHis and TEV protease cleavage site optimized for expression in *E. coli* was synthesized by GeneArt and subcloned into expression plasmid pETDuet-1. Protein expression was carried out in *E. coli* strain BL21 in TB media. After reaching $OD_{600}$=1.2 the bacteria were induced by adding 0.5 mM IPTG, cultured for 3 h at 37°C, and harvested by centrifugation. Cells were then resuspended in a lysis buffer containing 50 mM Tris pH 8.5, 300 mM NaCl, 5% glycerol, 1% Triton X-100, 2 mM β-mercaptoethanol, and protease inhibitor cocktail and disintegrated by sonication. The lysate was cleared by centrifugation, filtered through a 0.45 μm filter and the protein was purified by nickel affinity (Ni-NTA Agarose, Jena Bioscience). TEV cleavage was carried out overnight in 50mM Tris pH 8.5, 250 mM NaCl, 5% Glycerol and 4 mM β-mercaptoethanol. TEV and uncut $M^{pro}$ were removed by a second NiNTA purification step. The protein was further purified by size exclusion chromatography (Superdex s75, GE Healthcare) in 50mM Tris pH 7.4, 150mM NaCl, 2mM β-mercaptoethanol.

**$PL^{pro}$ activity assay**

The assay was designed to measure $PL^{pro}$ protease activity under screening conditions in white 384-well Optiplates. The assay buffer contained 50 mM Tris-HCl (pH 8.0), 0.01 % (w/v) BSA and 10 mM DTT. RLRGG-AMC was used as a fluorogenic substrate for $PL^{pro}$ 40 μl of $PL^{pro}$ protein (final concentration 60 nM) was incubated with 10 μl RLRGG-AMC substrate (final concentration 400 nM). The mixture (final volume 50 μl) was incubated for 30 min. The release of AMC ($\lambda_{ex}$ = 360 nm; $\lambda_{em}$ = 487 nm) was measured on an Envision plate reader (Perkin Elmer, Waltham, MA).

**$M^{pro}$ activity assay**

The main protease activity assay was adapted from previously described kinetic assays(*3*). The cleavage of the fluorogenic substrate MCA-AVLQSGFR-Lys(Dnp)-Lys-NH2 (Peptide Specialty Laboratories, Heidelberg) was monitored over 30 min using a microplate reader (PerkinElmer) with excitation and emission wavelengths of 320 nm and 405 nm, respectively. Different concentrations of acriflavine were incubated with 2 µM main protease before the reaction was initiated by the addition of fluorogenic substrate. The measurement was performed in Greiner black flat bottom 384-well microplates. The final reaction volume of 50 µl contained 2 µM main protease, 15 µM fluorogenic substrate and acriflavine in a buffer consisting of 50 mM Tris-HCl (pH 7.3), 150mM NaCl, 1 mM EDTA and 1% DMSO.

**Compound Screening**

40 µl of $PL^{pro}$ protein (final concentration 60nM) in assay buffer (50 mM Tris-HCl (pH 8.0), 0.01 % (w/v) BSA, and 10 mM DTT) was added to the screening plates using the MultiFlo dispensing system (BioTek). Compounds and DMSO (control sample) were directly pipetted into the screening plates to achieve a final concentration of 10 µM using a Sciclone G3 liquid handling workstation (PerkinElmer, Waltham, USA). The drug repurposing collection (*4*) was used to identify $PL^{pro}$ inhibitors. The controls in the plates (first two and last two columns) included DMSO control (negative control with no compound) and control without protein (to obtain the assay window). After addition of 10 µl RLRGG-AMC substrate (final concentration 400 nM) with the MultiFlo, AMC fluorescence ($\lambda_{ex}$ = 360 nm; $\lambda_{em}$ = 487 nm) was measured using a Envision plate reader immediately after substrate addition (time point 0) and after 30

min (time point 1). The average *Z′* factor for all plates was above 0.8, showing excellent screening quality. As a cut-off value for active compounds, we choose 4x the standard deviation below median of all compound-treated wells (calculated per plate). After removing compounds that generated high fluorescence signal at time point 0 (suggesting unspecific compound interference).

We also assessed $PL^{pro}$ inhibition with ACF and two derivatives of ACF, namely acridine orange base and acridine-3,6-diamine sulfate (**Supplementary Figure S1D**), as well as the previously published $PL^{pro}$ inhibitor GRL-0167 (*16*) (**Supplementary Figure S1E**). All derivatives showed dose-dependent inhibition of $PL^{pro}$, but were less active than the parent compound.

### $PL^{pro}$ inhibitor $IC_{50}$ determination

RLRGG-AMC or ISG15-AMC were used as substrates for $PL^{pro,}$ and the release of AMC fluorescence was measured ($\lambda_{ex}$ = 360 nm; $\lambda_{em}$ = 487 nm) on an Envision plate reader. 40 µl of a 75 nM $PL^{pro}$ solution in assay buffer (50 mM Tris-HCl (pH 8.0), 0.01% (w/v) BSA and 10 mM DTT) was pipetted into 384 well plates and different concentrations of ACF (50 µM – 0 µM, final concentration). The mixture was incubated for 1 h at RT. Then the reaction was initiated by adding 10 µl of 2 µM RLRGG-AMC (final concentration 400 nM) or 10 µl of 0.5 µM ISG15-AMC (final concentration 100 nM), respectively. Initial velocities of AMC release were normalized to the DMSO control. The $IC_{50}$ value was calculated using GraphPad Prism. The experiment was repeated three times.

### Gel-based $PL^{pro}$ DUB activity

0.2 µM $PL^{pro}$ was preincubated with 0, 10, 30 or 50 µM ACF for 30 min in 20 µl reaction buffer (20 mM Tris-HCl pH 7.5, 100 mM NaCl, 10 mM DTT) at 23°C. In the next step 20 µl of 4 µM tri-ubiquitin K48-linked (Boston Biochem) diluted in the reaction buffer was added to the pre-mix. Final enzyme and substrate concentrations were 0.1 µM and 2 µM, respectively. At indicated time points, the reaction was stopped by mixing 5 µl of the reaction with 15 µl 4x concentrated gel-loading buffer (Carl Roth), and the samples were analyzed by SDS–PAGE (Invitrogen NuPAGE™ 4–12% Bis-Tris) and western blot (anti-ubiquitin, P4D1, Santa Cruz Biotechnology).

#### Chemical characterization of the ACF components

The commercial ACF (Sigma-Aldrich/Merck #A8251) is sold as a mixture of 3,6-diaminoadridin-10-ium (proflavine) and 3,6-diamino-10-methylacridin-10-ium (acriflavine). In order to perform the study using fully characterized material, we have used NMR and HPLC/MS techniques to analyze the commercial product. The analysis revealed 56% proflavine (**PF**), 17% acriflavine (**ACF**), and 22-23% side methylated **PF**. The remaining 4-5% are other 3,6-diaminoacridine derivatives like for example, side methylated **ACF** (4%) (**See supplementary analysis**).

#### Cells and viruses

Vero (*Cercopithecus aethiops*; kidney epithelial; ATCC CCL-81), HCT-8 (ATCC CCL-244) cells, derivative of HRT-18 (ileocecal colorectal adenocarcinoma; ATCC CCL-244), CRFK (*Felis catus*, kidney cortex; ATCC® CCL-94) were cultured in Dulbecco's MEM (Thermo Fisher Scientific, Poland) supplemented with 5% fetal bovine serum (heat-inactivated; Thermo Fisher Scientific, Poland) and antibiotics: penicillin (100 U/ml), streptomycin (100 µg/ml), and ciprofloxacin (5 µg/ml). A549 cells with ACE2 overexpression (A549$^{ACE2+}$) (*8*) were cultured in the same manner but supplemented with G418 (5 mg/ml; BioShop, Canada).

LLC-MK2 cells (ATCC CCL-7; Macaca mulatta kidney epithelial cells) were maintained in minimal essential medium (MEM; two parts Hanks' MEM and one part Earle's MEM [Life Technologies, Poland]) 5% fetal bovine serum (heat-inactivated; Thermo Fisher Scientific, Poland), penicillin (100 U/ml), streptomycin (100 U/ml ), and ciprofloxacin (5 µg/ml).

Primary human skin fibroblasts (HSF) were cultured in Dulbecco's MEM (Thermo Fisher Scientific, Poland) supplemented with 10% fetal bovine serum (heat-inactivated; Thermo Fisher Scientific, Poland), 1% nonessential amino acids (Life Technologies) and antibiotics: penicillin (100 U/ml), streptomycin (100 µg/ml), and ciprofloxacin (5 µg/ml).

Human airway epithelial (HAE) cells were isolated from conductive airways resected from transplant patients. The study was approved by the Bioethical Committee of the Medical University of Silesia in Katowice, Poland (approval no: KNW/0022/KB1/17/10 dated 16.02.2010). Written consent was obtained from all patients. Cells were dislodged by protease treatment and later mechanically detached from the connective tissue. Further, cells were trypsinized and transferred onto permeable Transwell insert supports (ϕ = 6.5 mm). Cell differentiation was stimulated by the media additives and removal of media from the apical side after the cells reached confluence. Finally, cells were cultured for 4-6 weeks to form well-differentiated, pseudostratified mucociliary epithelium. All experiments were performed in accordance with relevant guidelines and regulations. Commercially available MucilAir™-Bronchial (Epithelix Sarl, Switzerland) HAE cultures were also used for the *ex vivo* analysis. MucilAir™ cultures were maintained as suggested by the provider in MucilAir™ culture medium. All cells were maintained at 37°C under 5% $CO_2$.

Reference SARS-CoV-2 strain 026V-03883 was kindly granted by Christian Drosten, Charité – Universitätsmedizin Berlin, Germany by the European Virus Archive - Global (EVAg); https://www.european-virus-archive.com/).

All SARS-CoV-2 stocks were generated by infecting monolayers of Vero cells. The cells were incubated at 37 °C under 5% $CO_2$. The virus-containing liquid was collected at day 2 post-infection (p.i.), aliquoted, and stored at −80°C. Control samples from mock-infected cells were prepared in the same manner.

MERS-CoV stock (isolate England 1, 1409231v, National Collection of Pathogenic Viruses, Public Health England, United Kingdom) was generated by infecting monolayers of Vero cells. The cells were incubated at 37°C under 5% $CO_2$. The virus-containing liquid was collected at day 3 p.i., aliquoted, and stored at −80°C. Control samples from mock-infected cells were prepared in the same manner.

FIPV stock (strain 79-1146) was generated by infecting CRFK cells at 90% confluency. The cells were incubated at 37 °C under 5% $CO_2$. The virus-containing liquid was collected at day 3 p.i., aliquoted, and stored at −80°C. Control samples from mock-infected cells were prepared in the same manner.

The HCoV-NL63 stock (isolate Amsterdam 1) was generated by infecting monolayers of LLC-MK2 cells. The cells were incubated at 32 °C under 5% $CO_2$ and then lysed by two freeze-thaw cycles at 6 days p.i. The virus-containing liquid was aliquoted and stored at −80°C. A control LLC-MK2 cell lysate from mock-infected cells was prepared in the same manner.

The HCoV-OC43 stock (ATCC: VR-1558) was generated by infecting monolayers of HRT-18 cells. The cells were incubated at 32 °C under 5% $CO_2$ and then lysed by two freeze-thaw cycles at 5 days post-infection (p.i.). The virus-containing liquid was aliquoted and stored at −80°C. A control HRT-18G cell lysate from mock-infected cells was prepared in the same manner.

Virus yields were assessed by titration on fully confluent cells in 96-well plates, according to the method of Reed and Muench (*9*). Plates were incubated at 32°C or 37°C, and the cytopathic effect (CPE) was scored by observation under an inverted microscope.

**Cell viability assay**

Cell viability was evaluated using the XTT Cell Viability Assay kit (Biological Industries, Cromwell, CT, USA) according to the manufacturer's protocol. Vero, A549$^{ACE2+}$, CRFK, HRT-18, LLC-MK2, and HSF cells were cultured on 96-well plates. Cells were incubated with ACF for 24 h at 37°C in an atmosphere containing 5% $CO_2$. After incubation, the medium was discarded and 100 μL of fresh medium was added to each well. Then, 25 μL of the activated 2,3-bis-(2-methoxy-4-nitro-5-sulphenyl)-(2H)-tetrazolium-5-carboxanilide (XTT) solution was added and samples were incubated for 2 h at 37°C. The absorbance (λ = 450 nm) was measured using a Spectra MAX 250 spectrophotometer (Molecular Devices, San Jose, CA, USA). The obtained results were normalized to the control samples, where cell viability was set to 100%.

**Virus replication inhibition assay**

Vero cells were seeded in culture medium on 96-well plates (TPP, Trasadingen, Switzerland) at 2 days before infection. Subconfluent cells were infected with SARS-CoV-2 viruses at 1600 50% tissue culture infectious dose ($TCID_{50}$)/ml. Infection was performed in the presence of 100 nM, 1 μM, and 10 μM concentrations of compounds listed in **Supplementary Table S1**. After 2 h of incubation at 37°C, cells were rinsed twice with PBS, and a fresh medium with compounds was added. The infection was carried out for 48 h, and the cytopathic effect (CPE) was assessed. Culture supernatants were collected from wells where CPE reduction was observed.

For the detailed determination of the antiviral properties of ACF, susceptible cells were seeded in a culture medium on 96-well plates at 2 days before infection. Subconfluent cells were infected with SARS-CoV-2, MERS-CoV and FIPV viruses at $TCID_{50}$ = 1600 and HCoV-

NL63 at $TCID_{50}$ = 4000 and HCoV-OC43 at $TCID_{50}$ = 3000. Infection was performed in the presence of ACF. Control cells were inoculated with the same volume of mock as a negative control and with 10 μM remdesivir as a positive control of coronavirus replication inhibition. After 2 h of incubation at 37°C, cells were rinsed twice with PBS, and fresh medium without or with ACF or remdesivir was added. The infection was carried out for 24 h when the CPE in virus control was observed. Culture supernatants were collected.

Virus replication inhibition *ex vivo* was evaluated by infecting HAE cultures with SARS-CoV-2 virus at 5000 $TCID_{50}$/ml in the presence of ACF, remdesivir or PBS. Two concentrations of ACF (400 nM and 500 nM) and the controls were added to the apical side of the inserts, followed by the addition of the virus diluted in PBS. Infection time was 2 hours at 32°C or 37°C. After the infection, the apical side of the HAEs were washed three times with PBS, and each compound was re-applied and incubated again for 30 minutes at 37°C. After the last incubation with the ACF, the samples (50 μl) were collected, and the HAEs were left in air-liquid interphase. Every 24 hours, the HAEs were incubated for 30 minutes with the ACF dilutions or controls, and the samples were collected. After collecting the last samples, cells were fixed with 3.7% paraformaldehyde and stained as described below. Virus yield was measured using the RT-qPCR method described below.

**Phylogenetic analysis**

The evolutionary history was inferred by using the Maximum Likelihood method and the General Time Reversible model (*10*). The tree with the highest log likelihood (-24114.75) is shown. Initial tree(s) for the heuristic search were obtained automatically by applying Neighbor-Join and BioNJ algorithms to a matrix of pairwise distances estimated using the Maximum Composite Likelihood (MCL) approach and then selecting the topology with a superior log-likelihood value. A discrete Gamma distribution was used to model evolutionary rate differences among sites (5 categories (+G, parameter = 2.6153)). The rate variation model allowed for some sites to be evolutionarily invariable ([+I], 4.79% sites). The tree is drawn to scale, with branch lengths measured in the number of substitutions per site. This analysis involved 17 nucleotide sequences. There were a total of 1576 positions in the final dataset. Evolutionary analyses were conducted in MEGA X (*11*).

**Isolation of nucleic acids, reverse transcription and quantitative PCR**

A viral DNA/RNA kit (A&A Biotechnology, Poland) was used for nucleic acid isolation from cell culture supernatants. RNA was isolated according to the manufacturer's instructions.

Viral RNA was quantified using quantitative PCR coupled with reverse transcription (RT-qPCR) (GoTaq Probe 1-Step RT-qPCR System, Promega, Poland) using CFX96 Touch real-time PCR detection system (Bio-Rad, Poland). The reaction was carried out in the presence of the probes and primers indicated in **Supplementary Table S2**. The heating scheme was as follows: 15 min at 45°C and 2 min at 95°C, followed by 40 cycles of 15 s at 95°C and 1 min at 58°C or 60°C. In order to assess the copy number of the N gene, standards were prepared and serially diluted.

**Virus visualization**

A549$^{ACE2+}$ cells were seeded on coverslips in 12 well plates (TPP, Trasadingen, Switzerland) at 2 days before infection. Subconfluent cells were infected with SARS-CoV---2 in the presence of ACF or remdesivir. After 2 h infection, unbound virions were washed off twice with PBS, and fresh medium supplemented with compounds was added. The infection was carried out for 24 h, and 48 h whereupon cells were fixed with 3.7% paraformaldehyde (PFA) for 1 h. Fixed cells were permeabilized using 0.5% Tween-20 (10 min, room temperature [RT]), and unspecific binding sites were blocked with 5% bovine serum albumin (BSA) in PBS

(4°C, overnight) prior to staining. For visualization of viral particles anti-SARS-CoV-2 nucleocapsid protein antibody (Bioss, bsm-41412M) at 1:200 dilution (2 h, RT) followed by Alexa Fluor 546 conjugated secondary antibody (Invitrogen, A-11003, 1:400 1 h, RT) was used. After incubating with antibodies, cells were washed thrice with 0.5% Tween-20. After labeling the virions, the actin cytoskeleton was visualized using Alexa-Fluor 647 conjugated Phalloidin (4 U/mL, 1 h, RT, Thermo Scientific), and nuclear DNA was stained with 4',6-diamidino-2-phenylindole (DAPI, 0.1 μg/ml, Sigma-Aldrich). Stained coverslips were mounted on glass slides with Prolong Diamond antifade mountant (Thermo Scientific, Poland). Fixed HAE cultures were permeabilized using 0.5% Triton X-100 (7 min RT), and unspecific binding sites were blocked with 5% BSA in PBS (1 h, 37°C) prior to staining. For visualization of virions anti-SARS-CoV-2 nucleocapsid protein antibody (Bioss bsm-41412M, 1:200 2 h, RT) coupled with goat anti-mouse Alexa Fluor 546 (Invitrogen, A-11003, 1:400 1 h, RT) were used. Nuclear DNA was stained with DAPI (0.1 μg/mL, 20 min, RT, Sigma-Aldrich) and actin cytoskeleton with Alexa Fluor 647 conjugated phalloidin (4 U/ml, 1 h, RT, Thermo Scientific). Stained HAE were cut from inserts and mounted on glass slides with cells facing coverslips. Fluorescent images were acquired under Zeiss LSM 880 confocal microscope.

**Animal study**

Transgenic mice expressing the human ACE2 protein under the human cytokeratin 18 promoter were purchased from the Jackson Laboratory. Mice were quarantined for at least 7 days prior to the experiment. Each experimental group consisted of 10 animals (14 animals in the control group). To take into account the fast metabolism of ACF, the compound was administered as a drinking solution (100 mg/kg body weight per day) and intramuscularly (5 mg/kg body weight or 15 mg/kg body weight twice a day). The treatment control group received remdesivir intramuscularly (25 mg/kg body weight per day). To mask the potentially unpleasant taste of the ACF that could lead to mice drinking less liquid than appropriate, all groups of mice were given sweetened isotonic drink instead of water. This is because, in our experience, the beverage is preferred by mice and well suited to dissolve the potentially unpleasant-tasting medications. The amount of drink consumed per day was determined prior to the experiment, and the concentration of ACF was adjusted to ensure the intended dose. Mice had free, permanent access to the drink during the experiment (from day -1 to 6 post-infection).

Mice treated intramuscularly received ACF every 12 hours as saline solution or remdesivir every 24 hours as a saline solution from day -1 until day 6 post-infection. No adverse effects were observed during the experiment.

On day 0 the animals were infected intranasally with the SARS-CoV-2 virus (Munchen-1.2 2020/984; 5 μl to each nostril) at 426,000 $TCID_{50}$/ml, which corresponded to ~$3\times10^5$ pfu. The virus was propagated and titrated on Vero cells. Infected mice were examined and weighed daily. On day 6 post-infection, animals were sacrificed under anesthesia. Lungs and brains were collected. Tissues were homogenized using a bead homogenizer (TissueLyser II, Qiagen, Poland). Viral RNA was isolated using mirVana™ miRNA Isolation kit (ThermoFisher Scientific, Poland), and the viral infection was quantified using the RT-qPCR.

### NMR spectroscopy

To assess the direct binding of ACF to $PL^{pro}$ in a solution, we produced $^{15}$N-labelled deuterated recombinant protein by growing the bacteria cultured in minimal M9 medium in $D_2O$. The protein purification is described earlier in the text. NMR experiments were recorded at 298 K on a Bruker Avance III 900 MHz spectrometer ($^1$H frequency 900 MHz) equipped with a 5 mm TCI cryoprobe. $^1$H,$^{15}$N TROSY spectra (*12*, *13*) were acquired with 40 scans and 1024 x 512 complex points on uniformly $^2$H,$^{15}$N-labelled $PL^{pro}$ at 170 μM concentration in PBS buffer, pH 7.4 supplemented with 10% $D_2O$. Spectra were processed and analyzed using the

AZARA suite of programs (v. 2.8, © 1993–2020; Wayne Boucher and Department of Biochemistry, University of Cambridge, unpublished). NMR titration experiments showed only localized alterations of spectra upon binding of ACF. No signs of oligomerization/aggregation were observed. This indicates that the third proflavine moiety observed in crystal structure at the interface between neighboring $PL^{pro}$ molecules is indeed a crystallization artifact and that ACF does not cause dimerization of the $PL^{pro}$ in solution. The multiple π-π stacked electron densities observed in the X-ray data between adjacent active sites also do not seem to cause $PL^{pro}$ oligomerization as this would cause many resonances to shift upon the addition of a high concentration of ACF.

**Crystallization and structure solution**

$PL^{pro}$ for crystallization was purified as described above. In the last step of purification, the protein was applied to size exclusion chromatography on S75 column using the following carrier buffer: 20 mM Tris-HCl pH 8.0, 40 mM NaCl, and 2 mM dithiothreitol (DTT). Purified protein was concentrated to 5-10 mg/ml using 30 kD cutoff Amicon Ultra Filter. The $PL^{pro}$-proflavine complex was prepared adding 5-fold molar excess of the compound to the concentrated protein. The final concentration of DMSO was kept below 5%. Crystallization samples were prepared using commercial kits and an automated crystallization workstation (Mosquito, TTP LabTech). Both ACF and pure proflavine were used in the experiments. Crystals of $PL^{pro}$-proflavine complex grew at room temperature in 0.05 M Hepes sodium salt pH 7, 0.05M magnesium sulfate, and 1.6 M lithium sulfate. Crystals suitable for testing were moved in cryo-protectant solution containing the harvesting solution supplemented with 25% (v/v) glycerol and snap-frozen in liquid nitrogen.

X-ray diffraction on $PL^{pro}$-proflavine crystals was determined at the Swiss Light Source (SLS, Villigen, Switzerland), beamline PXIII. The best dataset was collected at 2.7 Å resolution and it was indexed and integrated using XDS software(*14*); scaled and merged using STARANISO web server (*15*). The crystal belongs to space group $P6_522$. Matthews coefficient analysis suggested the presence of two $PL^{pro}$-proflavine molecules in the asymmetric unit. Molecular replacement solution was found using Phaser(*16, 17*) and the apo $PL^{pro}$ structure (PDB code: 6W9C) as a searching model. Model and restraints for proflavine were prepared using Lidia, the ligand builder in Coot(*18*). The initial model was subjected to several iterations of manual and automated refinement cycles using COOT and REFMAC5, respectively(*19, 20*). Throughout the refinement, 5% of the reflections were used for cross-validation analysis(*21*), and the behavior of $R_{free}$ was employed to monitor the refinement strategy. Detailed information on data collection and refinement statistics is listed in **Supplementary Table S3**. There are two molecules of the complex in the asymmetric unit. Intriguingly, a disulfide bond was found, which bridges two Cys270 of adjacent $PL^{pro}$ molecules from different asymmetric units. An additional proflavine molecule is located at the interface between these symmetry mates between two molecules and is considered a crystallographic artifact (**Supplementary Figure S7**). The electron density analysis shows a weaker trace of at least two more proflavines that can be allocated on top of the other two at optimal distance for $\pi$-$\pi$ staking forming a continuous, DNA-like stacking from one to another $PL^{pro}$ molecule in the same asymmetric unit. Their electron density is much weaker and does not allow these molecules to be to built unambiguously (**Supplementary Figure S8**). However, they are not involved in any interaction with $PL^{pro}$.

**Statistical analysis**

All experiments were performed in triplicate, and results are presented as mean ± SEM unless otherwise indicated. One-way ANOVA with Tukey HSD post-hoc test was used to assess statistical significance of obtained results. When the parametric test assumptions were violated, the nonparametric Kruskal-Wallis with Dunn's post-hoc test was used. P values of 0.05 and less were considered significant.
