## Supplementary Figures for "Acriflavine, a clinically aproved drug, inhibits SARS-CoV-2 and other betacoronaviruses"

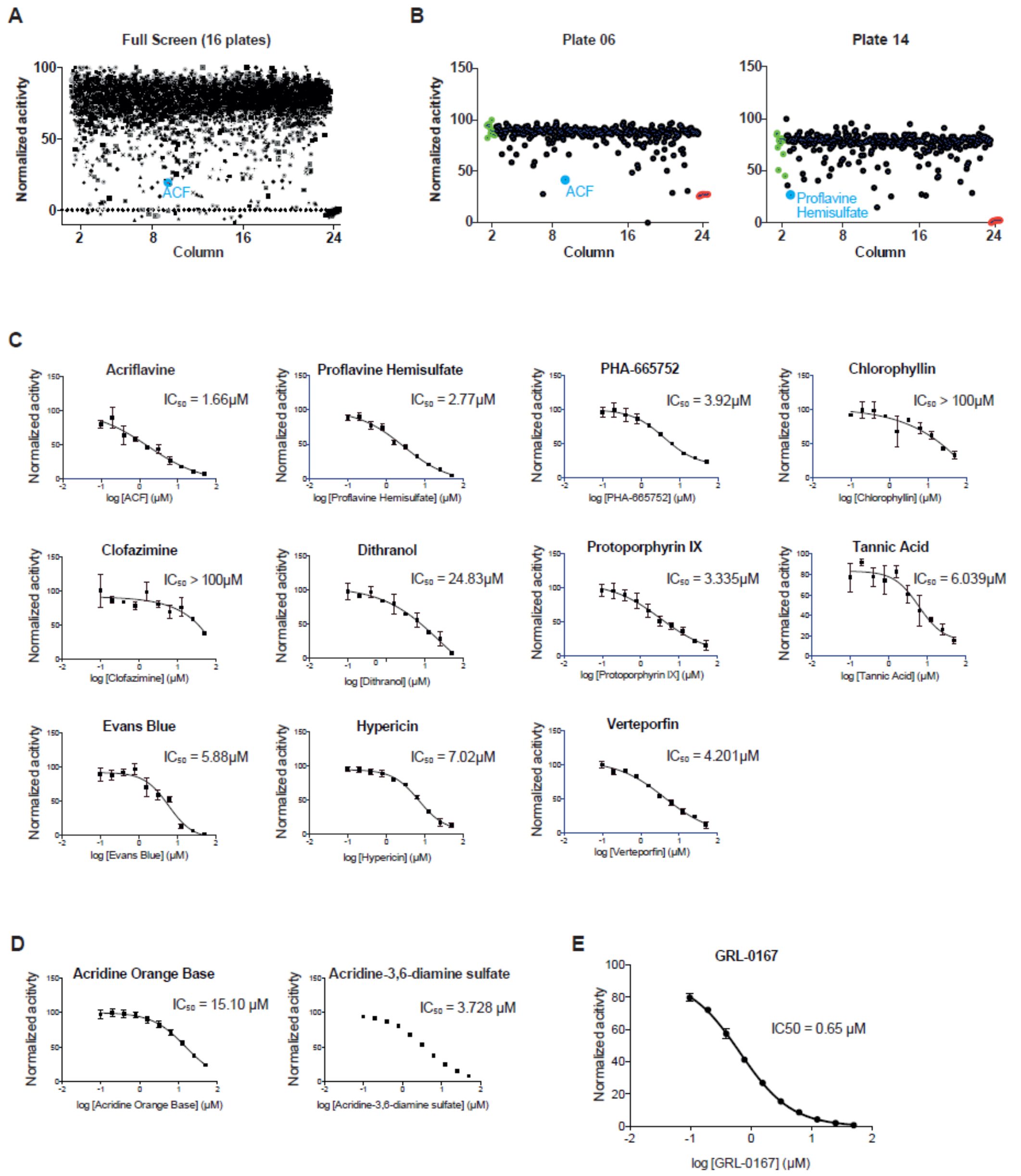


**Supplementary Figure S1.** (A) Summary of the high-throughput screening campaign (16 x 384-well plates). RLRGG-AMC peptide was used as a substrate for His-PL$^{pro}$. Each dot represents the data of one compound in one well (n = 1). Controls without protease are located in column 24, and ACF as a representative hit is depicted in blue. (B) Two representative screening plates with ACF and Proflavine Hemisulfate as hits (blue) are depicted side by side. (C) Dose-response curves of eleven hits. These hits were re-ordered and re-tested in ten-point titrations on the primary screening assay (mean ± SD, n = 2). Shown are inhibition curves and $IC_{50}$ values. (D) as in (C). Analogs of ACF were tested in ten-point titrations on the primary assay (mean ± SD, n = 3). (E) Inhibition curve of the known PL$^{pro}$ inhibitor GLR-0617 (mean ± SD, n = 3).

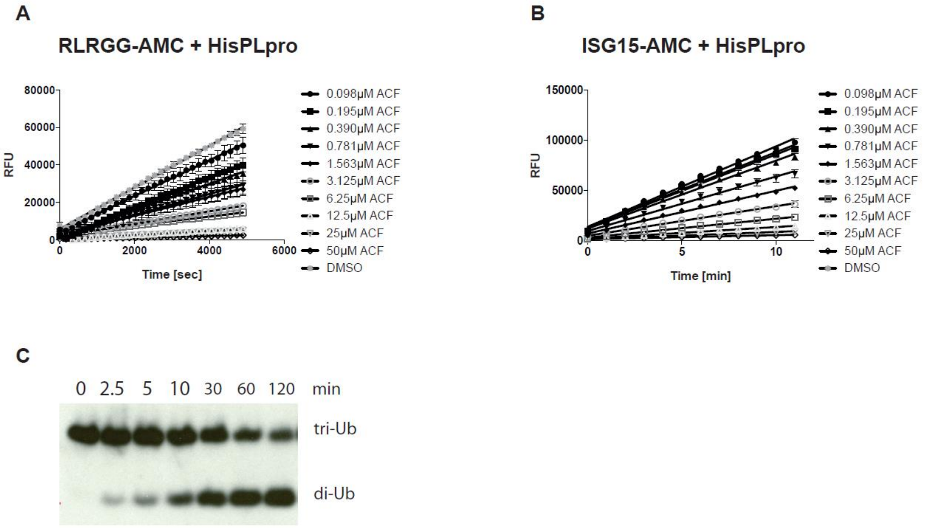


**Supplementary Figure S2.** (A) and (B) Kinetic assays of His-PL$^{pro}$ activity in the presence of different concentrations of ACF. RLRGG-AMC in (A) and ISG15-AMC in (B) were used as substrates. Reactions were performed in triplicates. (C) Time-course analysis of tri-ubiquitin K48-linked (2 µM) hydrolysis using 100 nM His-PL$^{pro}$.

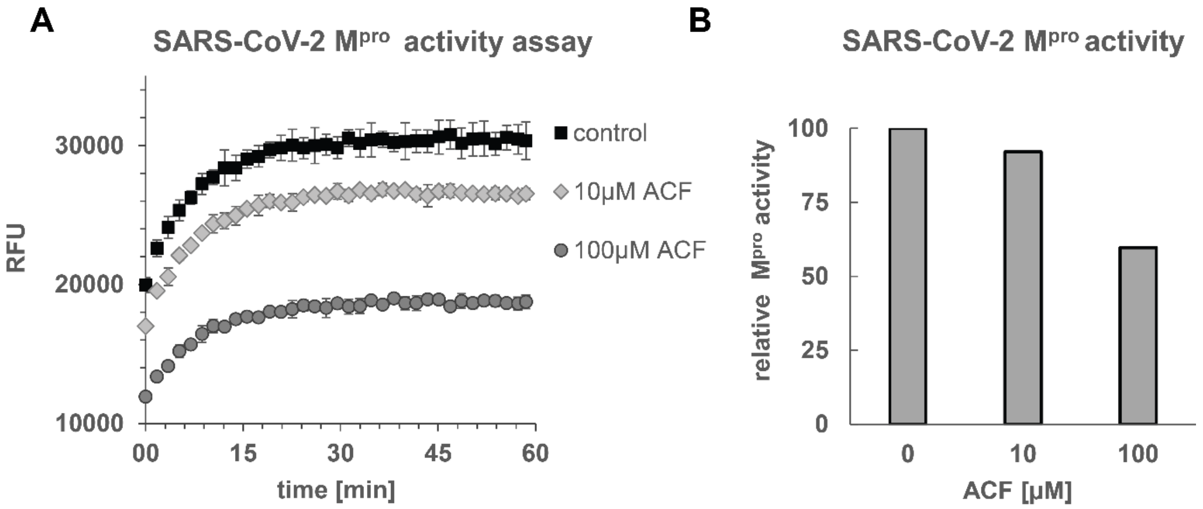


**Supplementary Figure S3.** SARS-CoV-2 $M^{pro}$ activity is not significantly inhibited by ACF. (A) The digestion of fluorogenic substrate was recorded in the absence and presence of ACF. The vertical shift in signal levels is caused by ACF absorbance. (B). Only a small decrease of $M^{pro}$ activity is observed at a physiologically irrelevant ACF concentration of 100 µM.

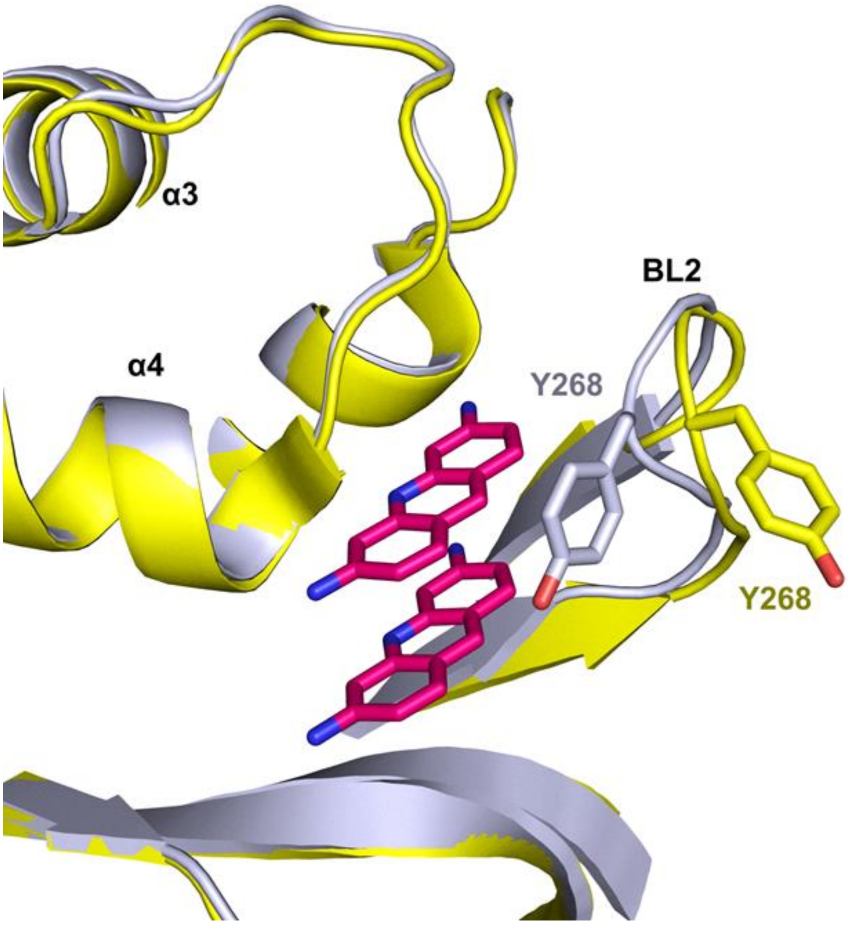


**Supplementary Figure S4.** Comparison of SARS-CoV2-PL$^{pro}$ in the proflavine-bound and apo-state. Bound PL$^{pro}$ is colored in gray; whereas the unbound PL$^{pro}$ (PDB ID: 7D47) is colored in yellow. The BL2 loop is involved in an induced-fit rearrangement upon the binding mostly due to the movement of the Tyr268. The side-chain of Tyr268 participates in a π-π stacking with proflavine molecules.

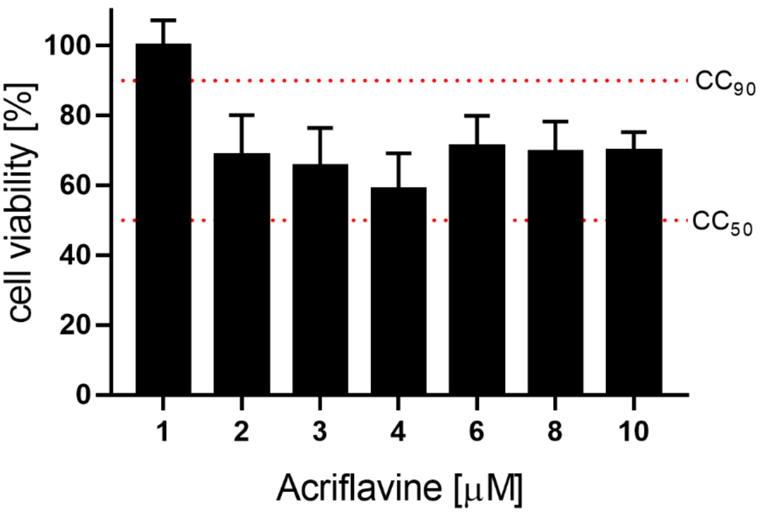


**Supplementary Figure S5.** The cytotoxicity of ACF in CRFK cells

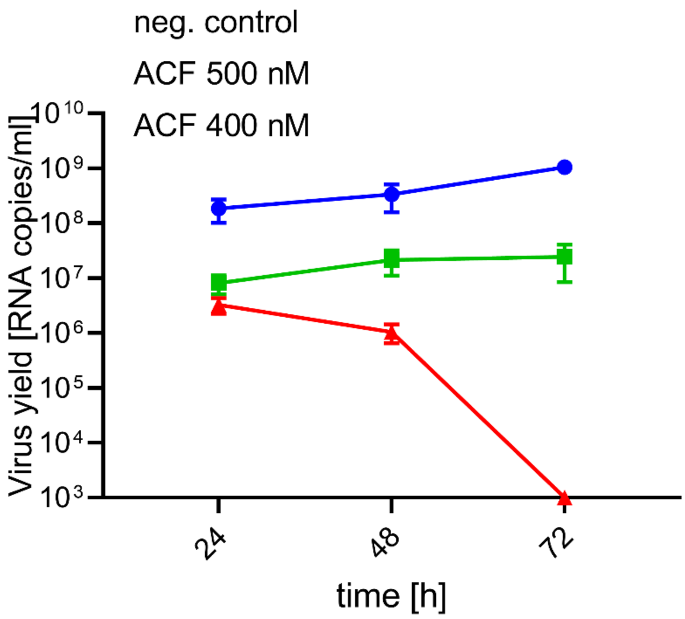


**Supplementary Figure S6. ACF blocks SARS-CoV-2 replication *in vitro* and *ex vivo*.** Antiviral activity of ACF against SARS-CoV-2 in human airway epithelium (prepared in-house). The figure shows RT-qPCR analysis of HAE culture supernatants infected with SARS-CoV-2. The assay was performed at least in duplicate, and median values with range are presented. Two-way ANOVA analysis with Dunnett's post-hoc test indicated that ACF and REM significantly inhibit virus yields during infection course compared to untreated control.

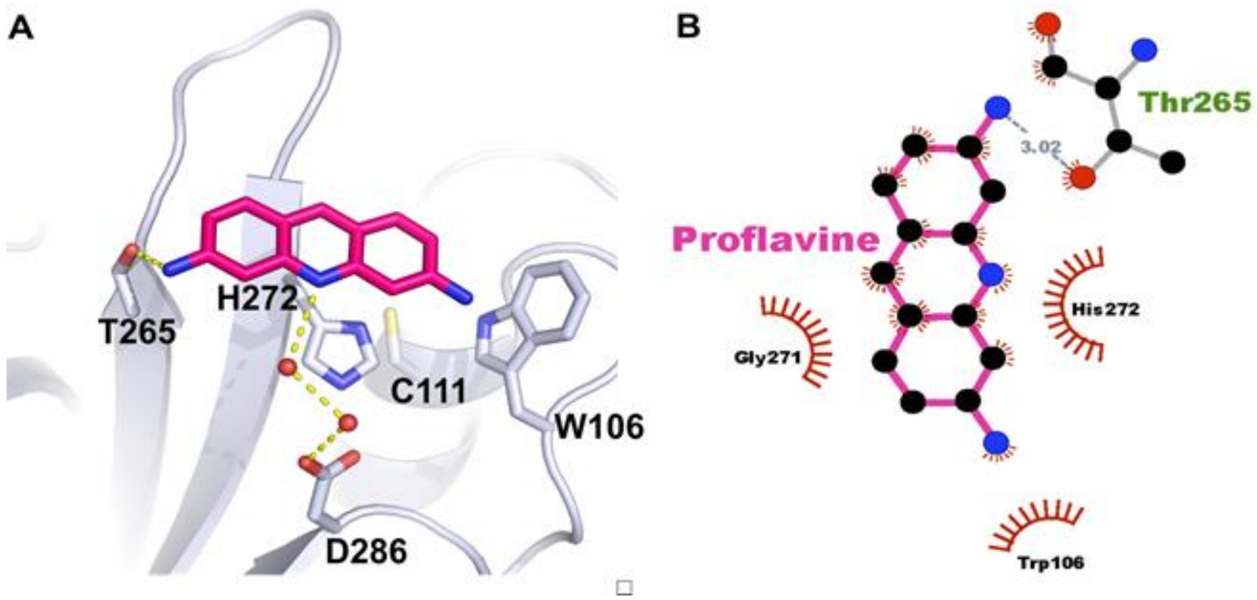


**Supplementary Figure S7.** Proflavine molecule at the interface between SARS-CoV2-PL$^{pro}$ asymmetric units. (A) Most probably due to a crystal packing molecule of proflavine was found on top of the catalytic triad (C111, H272, D286). Important residues are highlighted as stick model. Two water molecules (red spheres) mediate a hydrogen bond with D286. Hydrogen bonds are represented as yellow dashed lines. (B) 2D plot of the molecular interactions between proflavine and the residues of SARS-CoV-2-PL$^{pro}$.

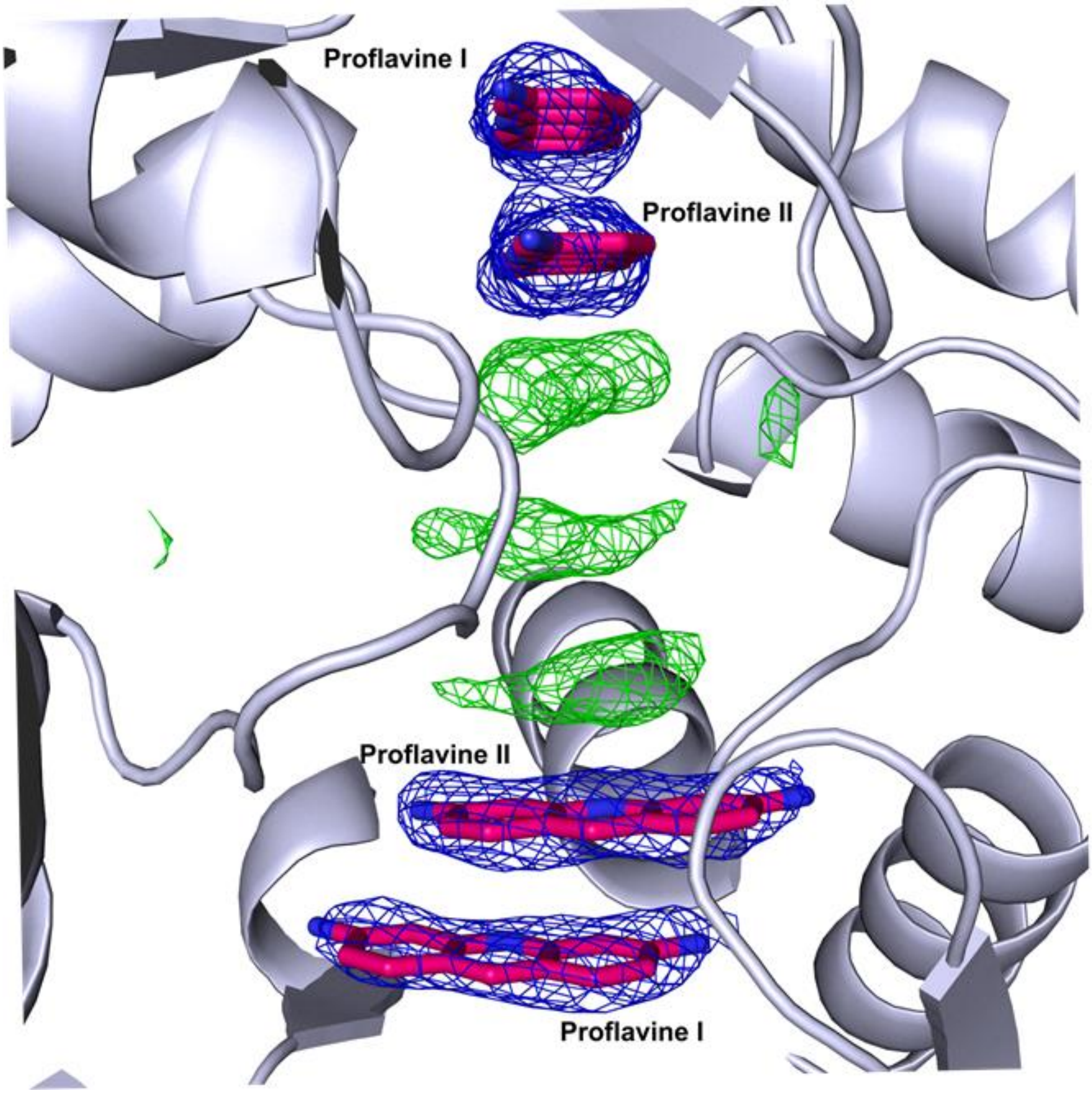


**Supplementary Figure S8.** Electron density map showing the fractional presence of additional aromatic proflavine-like molecules π-π stacked one on top of the other between two copies of SARS-CoV2-PL$^{pro}$ present in the crystal lattice. 2$F_o$-$F_c$ electron density map is contoured at 2σ. The electron density of the identified proflavine molecules is colored in blue; whereas additional electron density is colored in green. The densities are most likely caused by weak and transiently-bound proflavines. We did not model them in the crystal structure as their electron density was much weaker than active site-bound molecules.
