## Supplementary Tables for "Acriflavine, a clinically aproved drug, inhibits SARS-CoV-2 and other betacoronaviruses"

**Supplementary Table S1.** Compounds selected from the library for validation as PL$^{pro}$ inhibitors.

| no | Compound |
|---|---|
| 1 | Verteporfin |
| 2 | Evans Blue |
| 3 | Clofazimine |
| 4 | Hypericin from *Hypericus perforatum* |
| 5 | Dithranol |
| 6 | Chlorophyllin sodium copper salt |
| 7 | Acriflavine hydrochloride |
| 8 | Protoporphyrin IX |
| 9 | Tannic acid |
| 10 | PHA-665752 hydrate |
| 11 | Proflavine hemisulfate salt hydrate |

**Supplementary Table S2.** Primers and probes used for a RT-qPCR.

| Target | Primer | Sequence | Concentration [nM] |
|---|---|---|---|
| SARS-CoV-2 | Forward | CAC ATT GGC ACC CGC AAT C | 600 |
| | Reverse | GAG GAA CGA GAA GAG GCT TG | 800 |
| | Probe | ACT TCC TCA AGG AAC AAC ATT GCC A (FAM / BHQ1) | 200 |
| MERS-CoV | Forward | GGG TGT ACC TCT TAA TGC CAA TTC | 500 |
| | Reverse | TCT GTC CTG TCT CCG CCA AT | 500 |
| | Probe | ACC CCT GCG CAA AAT GCT GGG (FAM / TAMRA) | 200 |
| HCoV-NL63 | Forward | AAA CCT CGT TGG AAG CGT GT | 500 |
| | Reverse | CTG TGG AAA ACC TTT GGC ATC | 500 |
| | Probe | TGT TAT TCA GTG CTT TGG TCC TCG TGA T (FAM / TAMRA) | 100 |
| FIPV | Forward | TCT CGT GGT CGG AAG AAT AAT G | 500 |
| | Reverse | GAA CAA GGT CTC TCG GAC ATA AA | 500 |
| | Probe | CCC ATT ACC CTC GAA CAA GGA TCT (FAM/BHQ1) | 200 |
| HCoV-OC43 | Forward | AGC AAC CAG GCT GAT GTC AAT ACC | 500 |
| | Reverse | AGC AGA CCT TCC TGA GCC TTC AAT | 500 |
| | Probe | TGA CAT TGT CGA TCG GGA CCC AAG TA (FAM/BHQ1) | 200 |

**Table S3 Data collection and refinement statistics**

| | SCov2-PLpro-Proflavine (PDB ID:7NT4) |
|---|---|
| **Data collection** | |
| Space group | $P6_522$ |
| Cell dimensions | |
| *a*, *b*, *c* (Å) | 111.37, 116.37, 253.57 |
| α, β, γ (°) | 90, 90, 120 |
| Resolution (Å) | 49.4(2.68) * |
| Wavelength (Å) | 1.00 |
| $R_{pim}$ | 8.6(46.1) |
| $I / \sigma I$ | 51.9(1.8) |
| Completeness (%) | 95.7(72.2) |
| Redundancy | 38.1 |
| Observed reflections | 801001 (29586) |
| Unique reflections | 21442 (1073) |
| **Refinement** | |
| Resolution (Å) | 40 (2.68) |
| No. reflections | 22325 |
| $R_{work}$ / $R_{free}$ | 19.3/ 26.4 |
| No. atoms | |
| Protein | 2 |
| Ligand/ion | 6 |
| Water | 193 |
| *B*-factors | |
| Protein | 43.9 |
| Ligand/ion | 61.33 |
| Water | 25.2 |
| R.m.s. deviations | |
| Bond lengths (Å) | 0.014 |
| Bond angles (°) | 2.16 |
| Ramachandran statistics (%) | |
| Most favored regions | 93.3 |
| Additionally allowed regions | 5.2 |
| Generously allowed regions | 1.5 |

*A single crystal was used for data collection and structure determination. *Values in parentheses are for highest-resolution shell.
