## Supplementary material for "Acriflavine, a clinically aproved drug, inhibits SARS-CoV-2 and other betacoronaviruses": 6. ACF supplementary analysis

### Supplementary analysis of commercial acriflavine

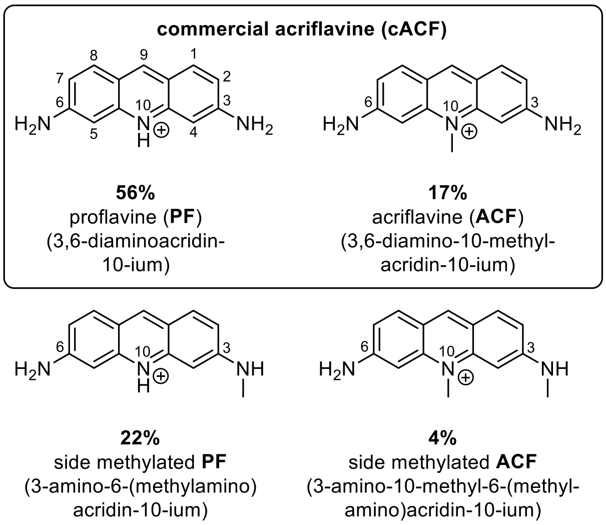


**Scheme S1.** Composition of commercial acriflavine (**cACF**) sold by Sigma-Aldrich/Merck (Catalogue number A8251): 56% proflavine (**PF**), 17% acriflavine (**ACF**) and about 26% of their side methylated derivatives.

All biological experiments were conducted using the same batch of commercial acriflavine hydrochloride (**cACF**) purchased from Sigma Aldrich (A8251), which is sold as a mixture of 3,6-diaminoadridin-10-ium (proflavine; **PF**) and 3,6-diamino-10-methylacridin-10-ium (acriflavine, **ACF**). In order to determine the exact composition of the obtained commercial acriflavine (**cACF**) the mixture was analyzed by HPLC-MS and $^{1}$H-NMR spectroscopy. Both methods show that the used batch of commercial acriflavine contains three main components and that the composition is as follows: 56% proflavine (**PF**), 17% acriflavine (**ACF**) and 22-23% side methylated **PF**. The remaining 4-5% are other 3,6-diaminoacridine derivatives like for example side methylated **ACF** (4%). The NMR analysis is based on the fact that the protons of the methyl group at position 10 of acriflavine (**ACF**) have a very distinct chemical shift of 3.94 ppm (4.04 ppm for side methylated **ACF**), whereas the protons of the methyl group connected to the amino group at position 3, like it is the case in side methylated **PF** and **ACF**, have a much lower chemical shift of 2.87 and 2.98 ppm, respectively.[1-3] The aromatic protons of the different 3,6-diaminoacridine derivatives almost exactly coincide with each other and consequently one of those peaks can be used for calibration of the integrals in the $^{1}$H-NMR spectrum. Integrating the overlapping doublets at 7.86-7.78 ppm as 2 protons and dividing the integrals of the separate signals for the methyl groups (4.04, 3.94 and 2.98 ppm) by 3 gives the percentages of side methylated **PF** (0.69:3 = 0.23; 23%) and side methylated **ACF** (0.13:3 = 0.04; 4%) as well as actual acriflavine (**ACF**) (0.52:3 = 0.17; 17%) present in the measured sample. As there are virtually no other 3,6-dimainoacridine derivatives other than **PF**, **ACF** and their side methylated derivatives, the remaining 56% can be attributed to proflavine (**PF**). This analysis fits perfectly the result obtained via the HPLC-MS technique. Commercial proflavine hemisulfate was also acquired from Sigma Aldrich and its purity was analyzed via HPLC-MS and $^{1}$H-NMR spectroscopy as well. The sample turned out to be 99% pure proflavine (**PF**) and was used as a reference for the analysis of commercial acriflavine (**cACF**). The signals found in the recorded $^{1}$H-NMR spectrum and the retention time obtained via the HPLC-UV-chromatogram (RT: 7.96 min) were in accordance with those of proflavine present in commercial acriflavine hydrochloride (RT: 7.89 min).

#### Experimental section

##### HPLC-UV/MS and $^{1}$H-NMR analyses of commercial acriflavine hydrochloride (cACF) and proflavine hemisulfate (PF)

HPLC-UV/MS analyses of commercial acriflavine hydrochloride and proflavine hemisulfate were performed on a Dionex UltiMate 3000 HPLC system coupled with a Thermo Finnigan LCQ ultrafleet mass spectrometer, using the following method: Waters X-Bridge C18 (4.6 x 30 mm, 3.5 µm) column; gradient: 5 to 10% of acetonitrile + 0.1% formic acid v/v in water + 0.1% formic acid v/v over 9.5 min perd; 10 to 20% of acetonitrile + 0.1% formic acid v/v in water + 0.1% formic acid v/v over 4.0 min period; 20 to 95% of acetonitrile + 0.1% formic acid v/v in water + 0.1% formic acid v/v over 1.0 min period; flow rate: 0.6 mL/min; UV detection at 254 nm. $^{1}$H-NMR spectra were recorded at room temperature on a Bruker AV-HD400 operating at 400 MHz.

[1]H-NMR spectrum of commercial acriflavine hydrochloride (**cACF**) recorded on a Bruker AV-HD400 (400 MHz, DMSO-d6):

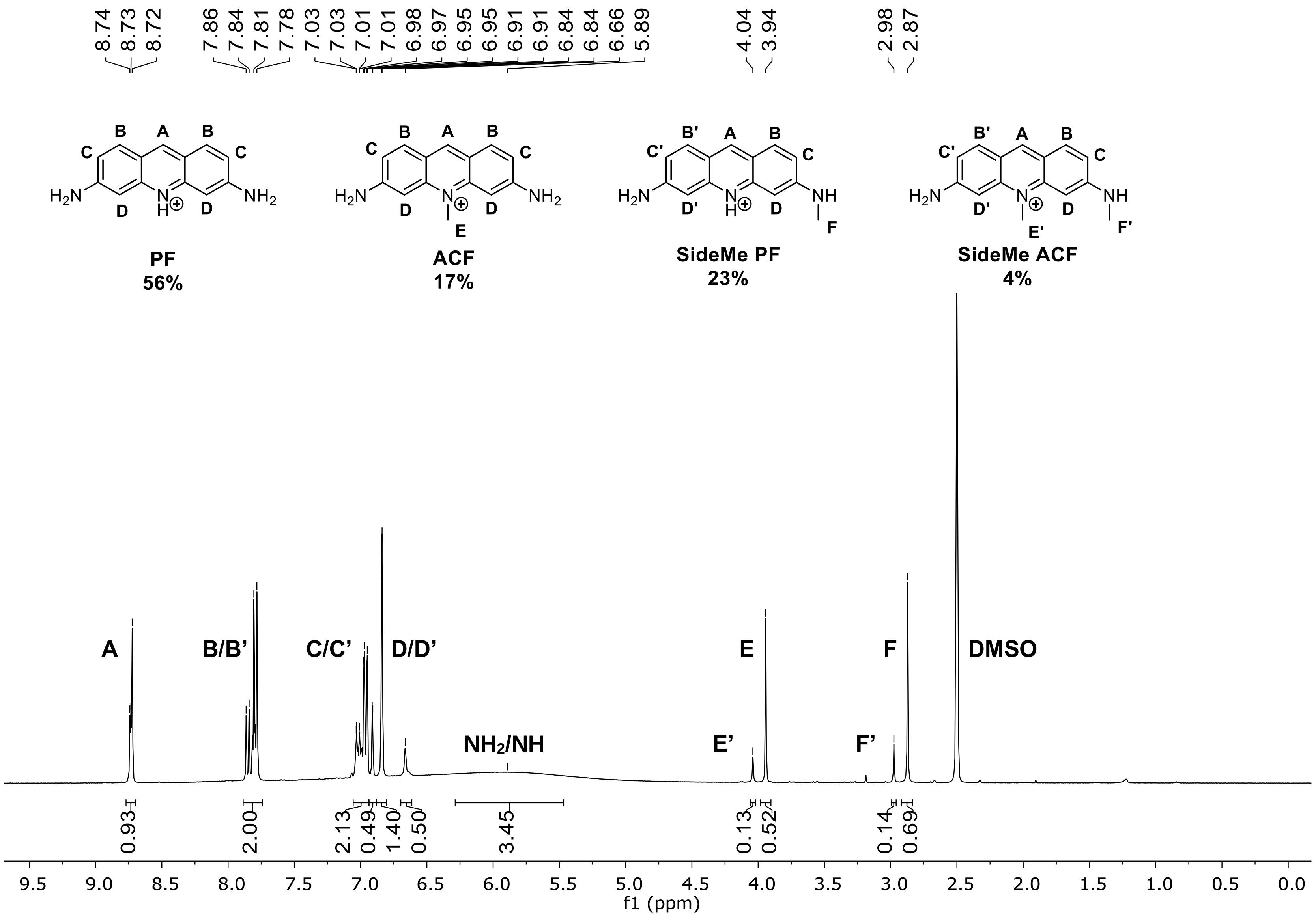

[1]H-NMR spectrum of commercial proflavine hemisulfate (**PF**) recorded on a Bruker AV-HD400 (400 MHz, DMSO-d6):

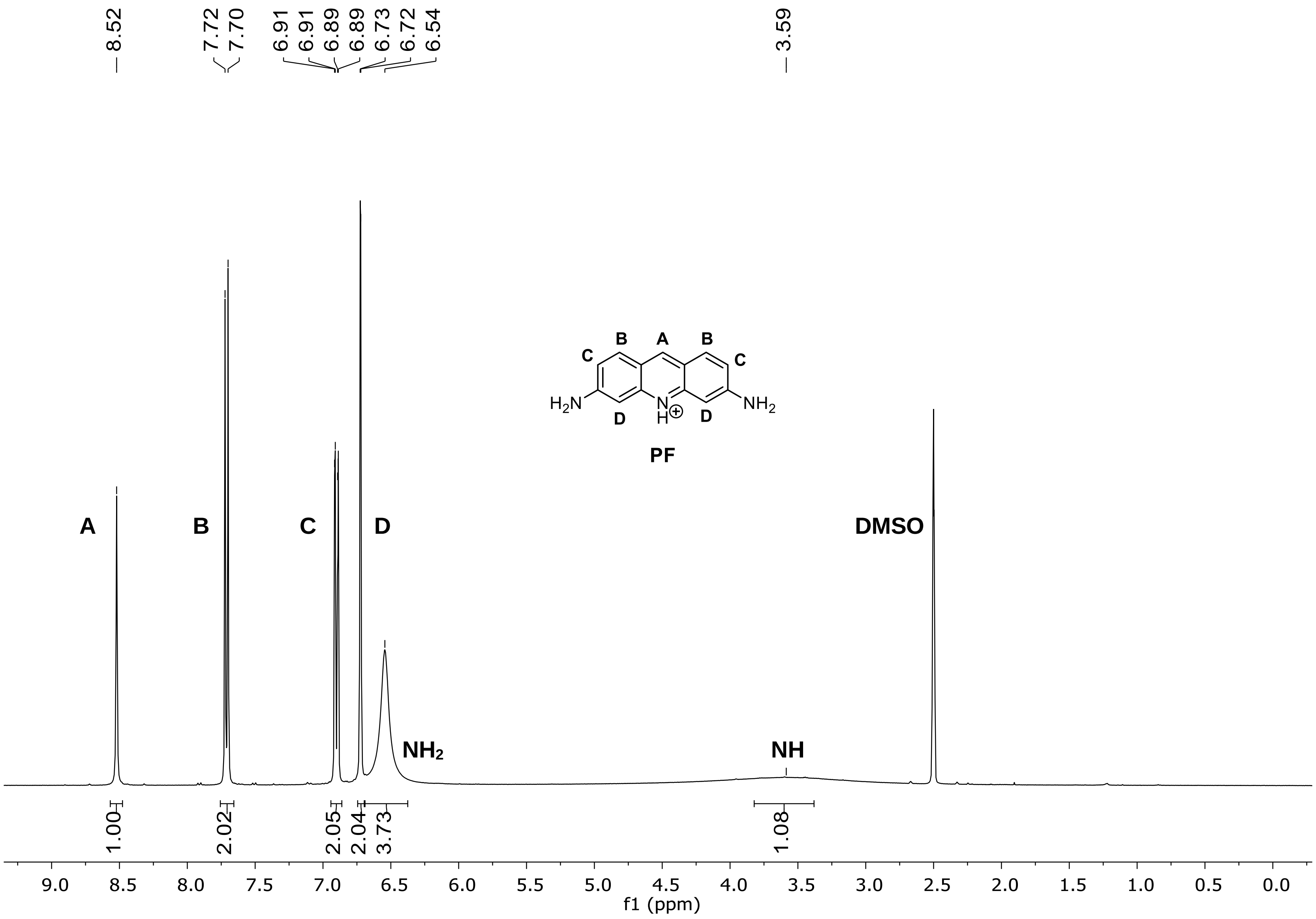

HPLC-UV-chromatogram and ESI mass spectra of commercial acriflavine hydrochloride (**cACF**) recorded on a Dionex UltiMate 3000 HPLC system coupled with a Thermo Finnigan LCQ ultrafleet mass spectrometer:

**Chromatogram**

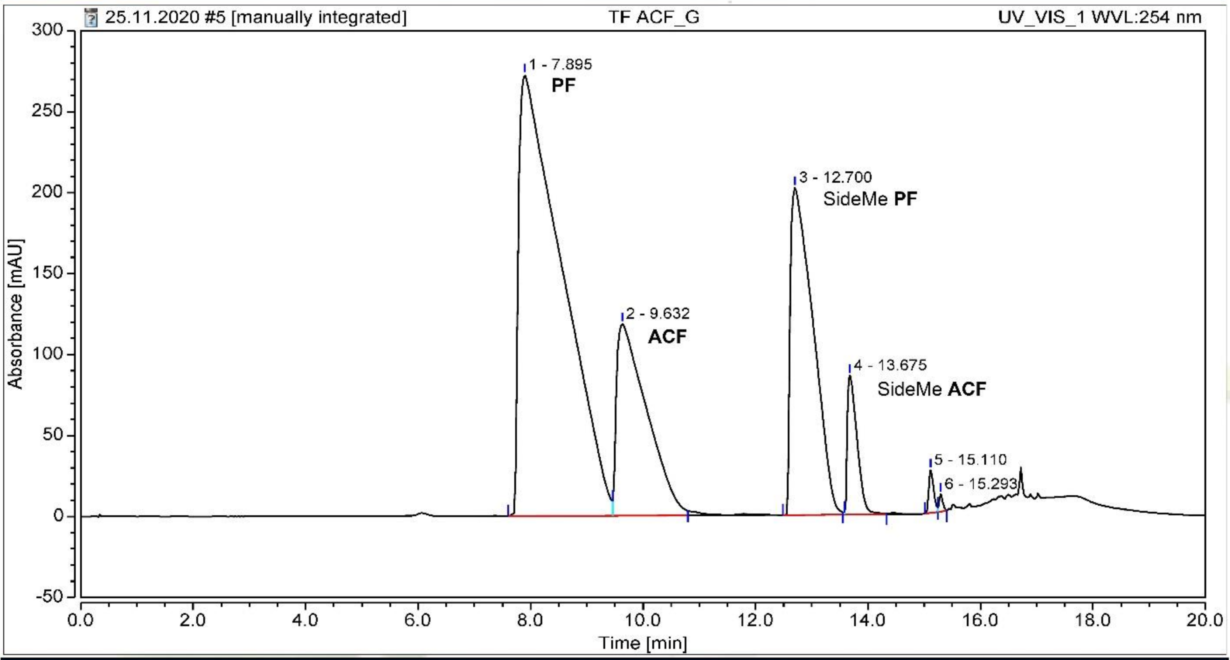


**Integration Results**

| No. | Peak Name | Retention Time min | Area mAU*min | Height mAU | Relative Area % | Relative Height % | Amount n.a. |
|---|---|---|---|---|---|---|---|
| 1 | Proflavine (**PF**) | 7.895 | 242.521 | 272.390 | 56.16 | 38.07 | n.a. |
| 2 | Acriflavine (**ACF**) | 9.632 | 73.680 | 118.343 | 17.06 | 16.54 | n.a. |
| 3 | SideMe **PF** | 12.700 | 94.706 | 202.156 | 21.93 | 28.25 | n.a. |
| 4 | SideMe **ACF** | 13.675 | 17.486 | 85.768 | 4.05 | 11.99 | n.a. |
| 5 | | 15.110 | 2.611 | 26.300 | 0.60 | 3.68 | n.a. |
| 6 | | 15.293 | 0.818 | 10.630 | 0.19 | 1.49 | n.a. |
| **Total:** | | | **431.823** | **715.588** | **100.00** | **100.00** | |

**Mass Spectrum**

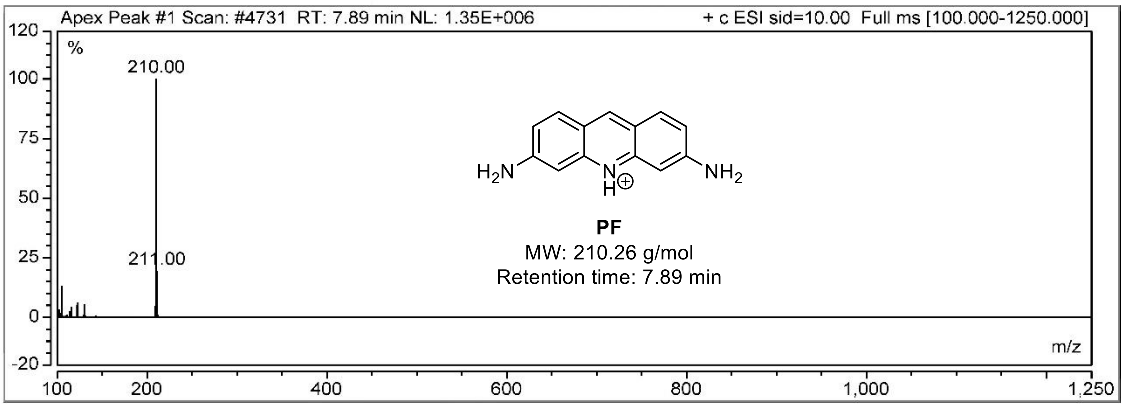

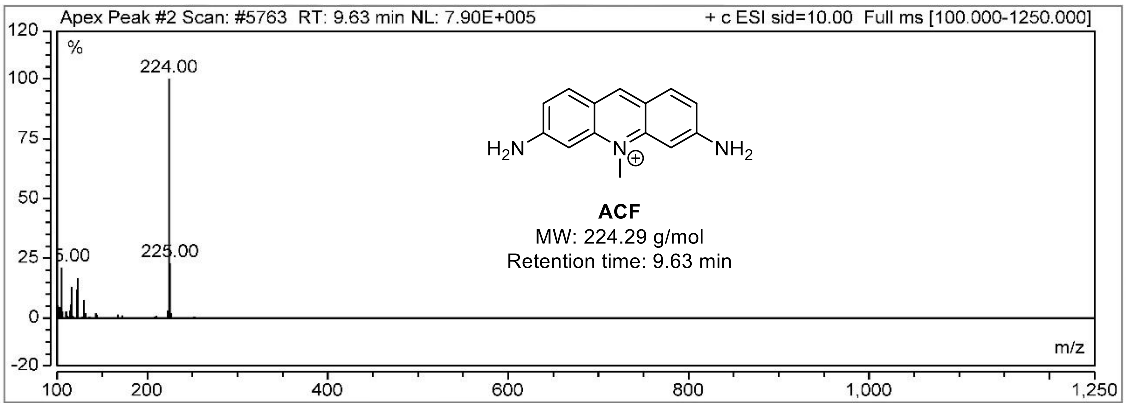
Apex Peak #2 Scan: #5763 RT: 9.63 min NL: 7.90E+005
+ c ESI sid=10.00 Full ms [100.000-1250.000]
%
224.00
225.00
m/z
ACF
MW: 224.29 g/mol
Retention time: 9.63 min

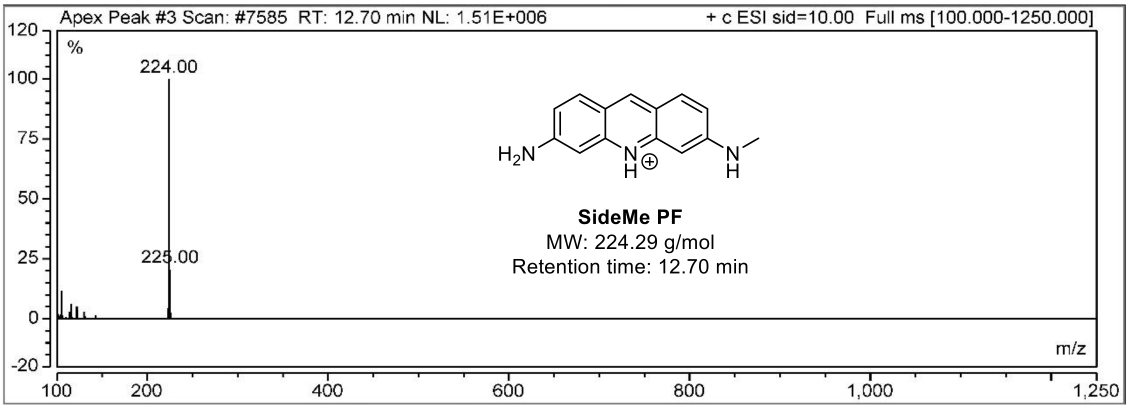
Apex Peak #3 Scan: #7585 RT: 12.70 min NL: 1.51E+006
+ c ESI sid=10.00 Full ms [100.000-1250.000]
%
224.00
225.00
m/z
SideMe PF
MW: 224.29 g/mol
Retention time: 12.70 min

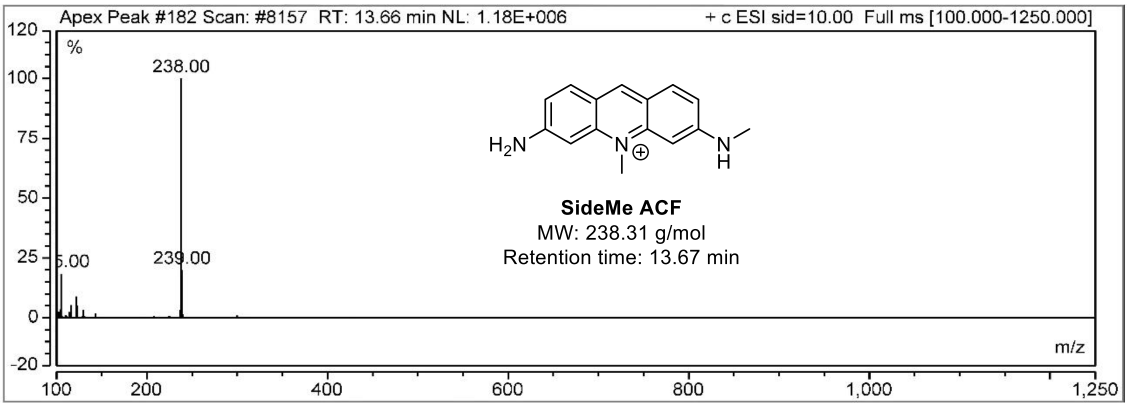
Apex Peak #182 Scan: #8157 RT: 13.66 min NL: 1.18E+006
+ c ESI sid=10.00 Full ms [100.000-1250.000]
%
238.00
239.00
m/z
SideMe ACF
MW: 238.31 g/mol
Retention time: 13.67 min

HPLC-UV-chromatogram and ESI mass spectra of commercial proflavine hemisulfate (**PF**) recorded on a Dionex UltiMate 3000 HPLC system coupled with a Thermo Finnigan LCQ ultrafleet mass spectrometer:

**Chromatogram**

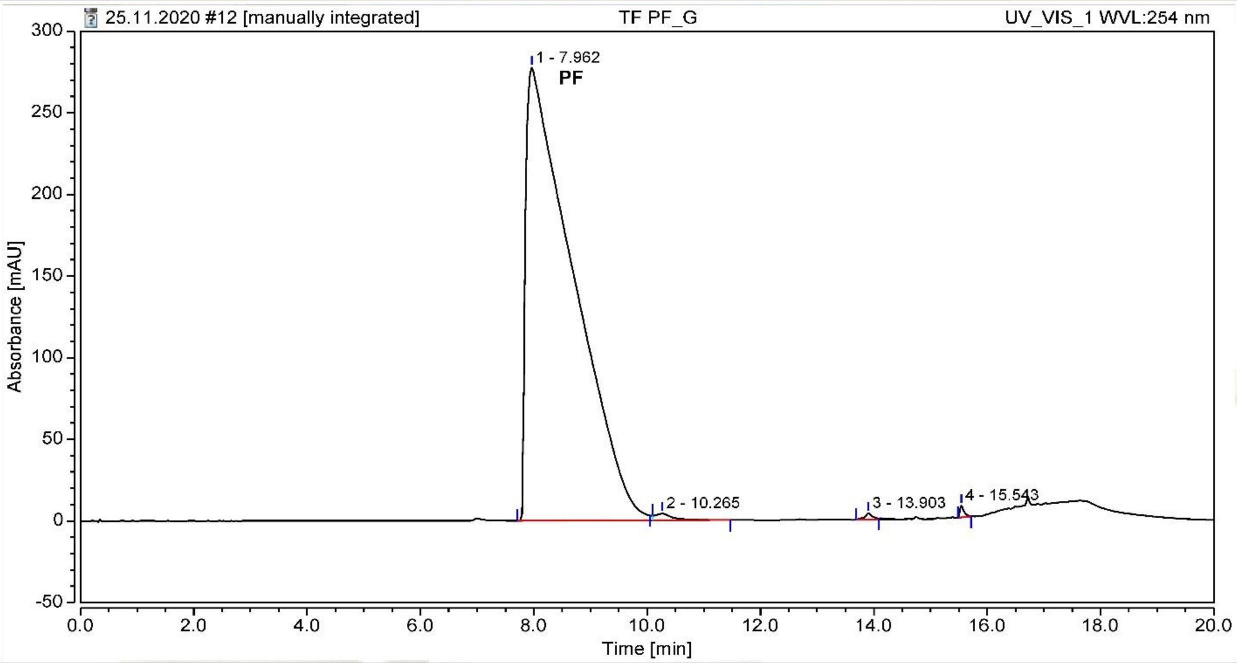


**Integration Results**

| No. | Peak Name | Retention Time min | Area mAU*min | Height mAU | Relative Area % | Relative Height % | Amount n.a. |
|---|---|---|---|---|---|---|---|
| 1 | Proflavine (**PF**) | 7.962 | 268.636 | 277.459 | 98.99 | 94.87 | n.a. |
| 2 | | 10.265 | 1.504 | 4.006 | 0.55 | 1.37 | n.a. |
| 3 | | 13.903 | 0.639 | 3.894 | 0.24 | 1.33 | n.a. |
| 4 | | 15.543 | 0.587 | 7.105 | 0.22 | 2.43 | n.a. |
| **Total:** | | | **271.366** | **292.464** | **100.00** | **100.00** | |

**Mass Spectrum**

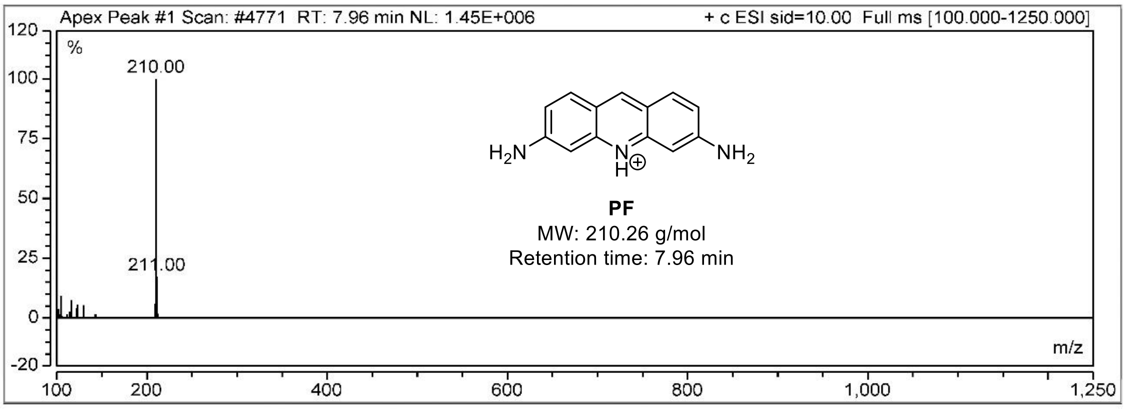
